# The ubiquitin-dependent ATPase p97 removes cytotoxic trapped PARP1 from chromatin

**DOI:** 10.1101/2021.07.16.452473

**Authors:** Dragomir B. Krastev, Shudong Li, Yilun Sun, Andrew Wicks, Daniel Weekes, Luned M. Badder, Eleanor G. Knight, Rebecca Marlow, Mercedes Pardo Calvo, Lu Yu, Tanaji T. Talele, Jiri Bartek, Jyoti Choudhary, Yves Pommier, Stephen J. Pettitt, Andrew Tutt, Kristijan Ramadan, Christopher J. Lord

## Abstract

Poly-(ADP-ribose) polymerase inhibitors (PARPi) elicit anti-tumour activity in homologous recombination defective cancers by promoting cytotoxic, chromatin-bound, “trapped” PARP1. How cells process trapped PARP1 remains unclear. By exploiting wild-type or trapping-resistant PARP1 transgenes combined with either a rapid immunoprecipitation mass-spectrometry of endogenous proteins (RIME)-based approach, or PARP1 Apex2-proximity labelling linked to mass-spectrometry, we generated proteomic profiles of trapped and non-trapped PARP1 complexes. This combined approach identified an interaction between trapped PARP1 and the ubiquitin-regulated p97 ATPase (aka VCP). Subsequent experiments demonstrated that upon trapping, PARP1 is SUMOylated by the SUMO-ligase PIAS4 and subsequently ubiquitinated by the SUMO-targeted E3-ubiquitin ligase, RNF4, events that promote p97 recruitment and p97 ATPase-mediated removal of trapped-PARP1 from chromatin. Consistent with this, small molecule p97 complex inhibitors, including a metabolite of the clinically-used drug disulfiram (CuET) that acts as a p97 sequestration agent, prolong PARP1 trapping and thus enhance PARPi-induced cytotoxicity in homologous recombination-defective tumour cells and patient-derived tumour organoids. Taken together, these results suggest that p97 ATPase plays a key role in the processing of trapped PARP1 from chromatin and the response of homologous recombination defective tumour cells to PARPi.

## Introduction

PARP inhibitors (PARPi) selectively kill tumour cells with impaired homologous recombination (HR). Based on this synthetic lethality, a number of PARPi have received regulatory approval for the treatment of breast, ovarian, pancreatic or prostate cancers with HR defects, including those with deleterious mutations in *BRCA1* or *BRCA2*^1^. The key target of PARPi, Poly(ADP-Ribose) Polymerase 1 (PARP1/ARTD1), is an ubiquitously-expressed nuclear enzyme that uses NAD^+^ as a substrate to synthesise linear and branched poly(ADP-ribose) (PAR) chains on substrate proteins (heteromodification) and itself (automodification). This catalytic activity (PARylation), which is activated by PARP1 binding to damaged DNA, initiates DNA repair by driving the recruitment of additional DNA repair proteins as well as by modulating chromatin structure. Once DNA repair is initiated, PARP1 is released from DNA via auto-PARylation. The majority of clinical PARPi bind the NAD^+^ binding site in the catalytic domain and inhibit PARP1 catalytic activity, but also induce the chromatin retention of PARP1 (PARP trapping), this latter characteristic being a significant driver of the cytotoxicity that PARPi elicit^2^. Consistent with the hypothesis that PARP1 trapping is a key determinant of PARPi-induced tumour cell cytotoxicity, deletion of *PARP1* causes PARPi resistance, as do in frame *PARP1* insertion/deletion mutations that impair PARP1 trapping^3^. Moreover, the chemical modification of a PARPi with poor trapping properties into a derivative compound with enhanced trapping property but similar catalytic potency, enhances cytotoxicity^4^. Although it is known that specific PARP1 mutations alter PARP1 trapping^3^, as does modulating the amount of residual PAR on PARP1 via the PAR-glycosylase (PARG)^5^, there is only a limited understanding of how trapped PARP1 is released from damaged DNA. This is one of the essential questions in PARP1 chromatin biology as mechanisms involved in removal of trapped PARP1 from chromatin represent potential modifiers of response to PARPi.

By generating a series of protein/protein interaction profiles of either trapped or non-trapped PARP1, we show here that trapped PARP1 binds the p97 ATPase, also known as Valosin Containing Protein (VCP). P97 is a hexameric unfoldase/segregase which uses ATP hydrolysis to unfold and disassemble ubiquitylated substrates through its central pore^6, 7^. It acts in various cellular locations including chromatin, on substrates such as Aurora B kinase, CMG helicases, the licensing factor CDT1, RNA pol II or the TOP1-cleavage complex^8–10^. We show that this interaction is mediated by sequential PIAS4-mediated SUMOylation and RNF4-mediated ubiquitylation that ultimately leads to the removal of trapped PARP1 from chromatin. In addition, we show that p97 inhibition, using a metabolite of the clinically used drug disulfiram, leads to prolonged PARP1 trapping and profound sensitisation to PARP inhibitors, suggesting an approach to enhancing PARPi-induced cytotoxicity. Collectively, our findings suggest that the p97-PARP1 axis is essential for removal of trapped PARP1 and the cellular response to PARPi.

## Results

### Identification of trapped PARP1-associated proteins

To understand the nature of the trapped PARP1 complex, we used two orthogonal systems to generate mass-spectrometry-based PARP1 protein/protein interactomes from cells with either trapped or non-trapped PARP1: (i) Rapid Immunoprecipitation Mass spectrometry of Endogenous proteins (RIME^11^) and (ii) *in vivo* Apex2 peroxidase-mediated labelling of proximal proteins^12^. To use these, we first generated cell lines derived from a previously described PARPi-resistant PARP1 defective cell line, CAL51 *PARP1^−/−^*^3^, into which we introduced one of three piggyBac transposon-based transgenes. For RIME profiling, we introduced either wild-type *PARP1* cDNA fused to an eGFP-coding sequence (*PARP1^WT^-eGFP*) or a *PARP1^del.p.119K120S^-eGFP* transgene. As we have previously shown that deletion of PARP1 residues p.119K120S (within the ZnF2-DNA binding domain) prevents PARP1 trapping by PARPi^3^, we used cells with the *PARP1^del.p.119K120S^-eGFP* transgene to allow RIME profiling of the PARP1 interactome in a setting where PARP1 could not be trapped (Figure 1A). To enable Apex2 profiling, we introduced into CAL51 *PARP1^−/−^* cells a wild-type *PARP1* cDNA transgene fused to Apex2 and eGFP coding sequences at the C-terminus (*PARP1^WT^-Apex2-eGFP*). After transposition and single-cell cloning, we established daughter clones that expressed the desired PARP1 fusion proteins (Supplementary Figure 1A). As expected, expression of either PARP1^WT^-eGFP *or PARP1*^WT^*-APEX2-GFP* proteins re-established PARPi sensitivity in otherwise resistant CAL51 *PARP1^−/−^* cells (Figure 1B, C), suggesting that it interacted with, and could be trapped by, a PARPi. Conversely, expression of the PARP1^del.p.119K120S^-eGFP protein did not re-establish PARPi sensitivity, consistent with the PARP1^del.p.119K120S^ mutant being trapping-defective^3^. Using a PAR-binding PBZ-mRuby2 probe and a UV microirradiation assay to estimate the extent of PAR production at UV-generated DNA damage sites^3^, we established that PARP1^WT^-Apex2-eGFP localised to DNA damage sites where it generated PAR (Figure 1D). In addition, PARP1^WT^-Apex2-eGFP could be “trapped” on damaged DNA by PARPi exposure (Figure 1D), exhibiting similar localisation kinetics to PARP1 protein fused to eGFP without the Apex2 tag (as described in^3^).

**Figure 1.**
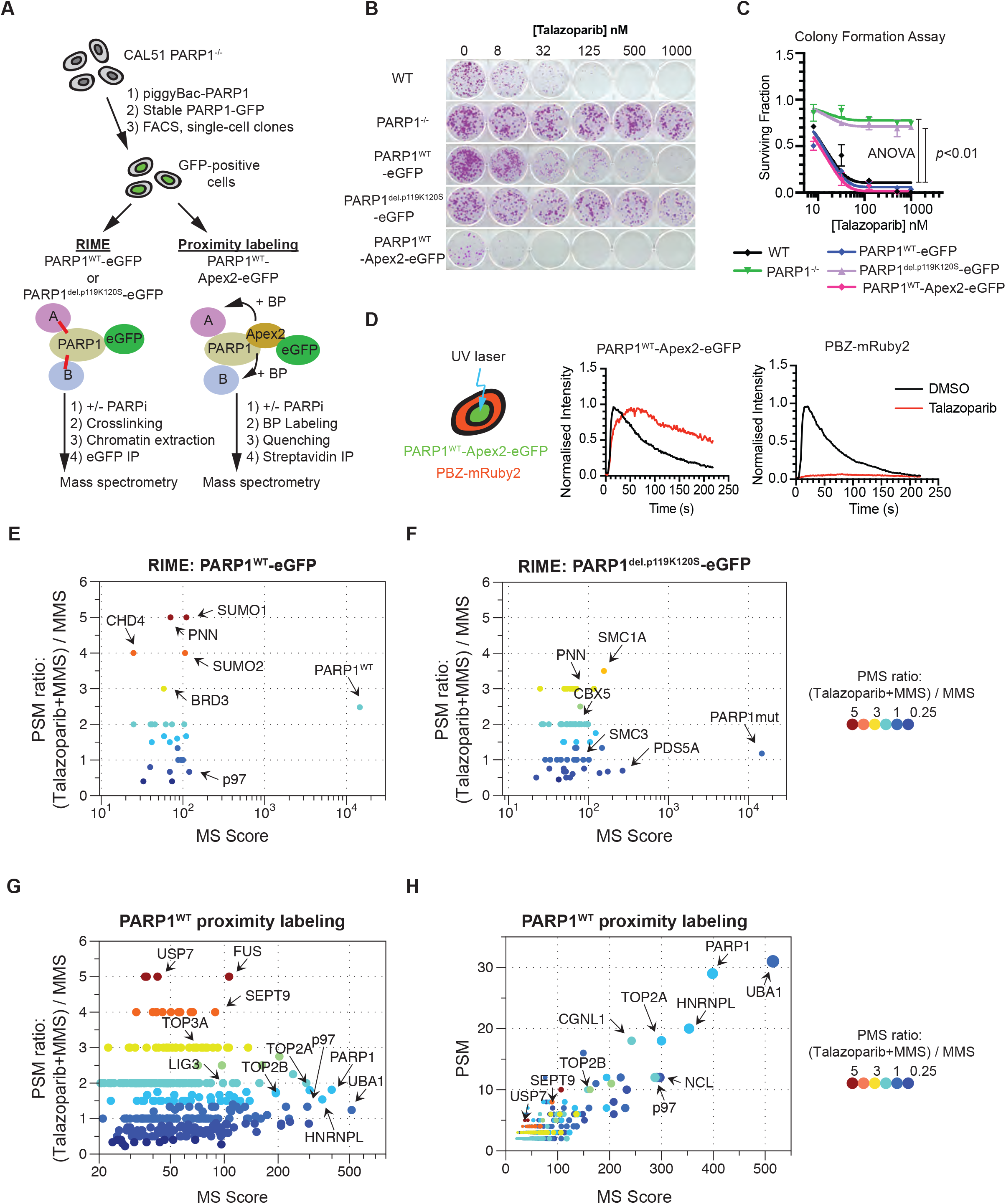
Identification of trapped PARP1 interacting proteins. **A.** Schematic describing the identification of trapped PARP1 protein/protein interactomes via RIME or proximity labelling linked to mass-spectrometry. CAL51 PARP1^−/−^ cells were transfected with a PARP1-encoding piggyBac transposons expressing either wild-type PARP1 fused to eGFP (PARP1^WT^-eGFP), a non-trapping mutant, PARP1^del.p.119K120S^-eGFP, or PARP1^WT^-Apex2-eGFP. After transposition (via a transposase-expressing plasmid), piggyBac positive cells were isolated by G418 selection and individual GFP-positive clones isolated by FACS. For the PARP1 RIME experiment (bottom left) *PARP1^−/−^*, PARP1^WT^-eGFP or PARP1^del.p.119K120S^-eGFP expressing cells were exposed to PARPi/MMS (to enable trapping) or MMS (no PARP1 trapping) for 1 hour, after which cells were cross-linked with formaldehyde. After quenching the reaction, the chromatin fraction was isolated and PARP1-associated proteins purified via GFP-Trap immunoprecipitation. Precipitates were then divided into aliquots for western blotting or on-bead tryptic digest followed by mass spectrometry analysis. A PARP1^WT^-Apex2-eGFP expressing clone was subjected to proximity labelling (bottom right) as follows: cells were exposed to PARPi/MMS (to enable PARP1 trapping) or MMS only (no PARP1 trapping) for 1 hour. In addition, the cells were exposed to biotin-phenol in the last 30 min of the treatment and then to H_2_O_2_ (BP labelling), followed by quenching of the labelling reaction. Biotinylated proteins were then purified by streptavidin immunoprecipitation (IP) under stringent conditions. Precipitates were then divided into aliquots for western blotting or on-bead tryptic digest followed by mass spectrometry analysis. **B.** Clonogenic assay illustrating PARP inhibitor sensitivity in CAL51 *PARP1^−/−^* cells expressing different PARP1 cDNAs. Cells described in (A) were exposed to talazoparib for 14 continuous days. Deletion of *PARP1* in CAL51 cells caused PARPi resistance; this resistance was reversed by expression of PARP1^WT^-eGFP or PARP1^WT^-Apex2-eGFP. Expression of a DNA-binding deficient form of PARP1, PARP1^del.p.119K120S^-eGFP, did not restore PARPi sensitivity. PARP1 protein expression in the different clones is shown in Supplementary Figure 1A. **C.** Quantification of colony formation assay shown in (B); ANOVA *p*-values are shown. **D.** PARP1^WT^-Apex2-eGFP protein localises to DNA damage, generates PAR and can be trapped by PARPi. Experimental schematic (left) -PARP1^WT^-Apex2-eGFP expressing cells were transfected with a PAR sensor, a PBZ PAR binding domain fused to mRuby2, which enables monitoring of PAR accumulation at sites of DNA damage caused by UV laser stripe. The two graphs illustrate the accumulation of PARP1^WT^-Apex2-eGFP-mediated fluorescence at the UV stripe over time (left) and the accumulation of PBZ-mRuby2 fluorescence (monitoring PAR) in the same experiment (right). Exposure of cells to 100 nM talazoparib, causes sustained accumulation of the GFP signal (i.e. PARP1^WT^-Apex2-eGFP trapping, left), but abolishes PAR production, as shown by the reduced PBZ-mRuby2 fluorescence (right). **E and F.** PARP1 interactions that are enriched under PARP1 trapping conditions (as defined by PSM ratio and MS scores – see Methods). Scatter plots are shown for PARP1^WT^-eGFP RIME (E) and PARP1^del.p.119K120S^-eGFP RIME (F). **G.** PARP1 interactions that are enriched under PARP1 trapping conditions for PARP1^WT^-Apex2-eGFP proximity labelling. **H.** A graph plotting the PSM against MS score for PARP1^WT^Apex2-eGFP proximity labelling interactions shows that p97 is among the most abundant proteins identified in the PARP1^WT^-Apex2-eGFP proximity labelling.

As PARP1 translocates to chromatin upon DNA damage, we first used RIME-based immunoprecipitation^11, 13^, to identify proteins associated with trapped PARP1 (Figure 1A). In these RIME experiments, protein interactions were first stabilised by crosslinking, after which chromatin-bound proteins were isolated. PARP1-associated complexes were then immunoprecipitated (via the eGFP tag) and subsequently identified by mass spectrometry. Cells expressing either PARP1^WT^-eGFP or PARP1^del.p.119K120S^-eGFP were first exposed to trapping conditions (MMS + talazoparib) after which, PARP1-interacting proteins identified by RIME. In this experiment, CAL51 *PARP1*^−/−^ cells were used as a control to allow proteins non-specifically bound to beads to be removed from the analysis (Supplementary Figure 1B). From this RIME analysis, we identified 50 proteins associated with PARP1 in cells expressing wild-type PARP1 either in the presence or absence of PARPi (Supplementary Table 1) and 144 PARP1-associated proteins in cells expressing the *PARP1^del.p.119K120S^-eGFP* transgene (Supplementary Table 2). In both datasets PARP1 was by far the most abundant protein identified by MS score (Figure 1E, F). In order to prioritise proteins for further analysis, we examined MS score (a combination score representing the peptide abundance and the quality of identification) and the enrichment ratio of peptide spectrum matches (PSM) in the methyl methanesulfonate (MMS)+talazoparib exposed cells compared to cells exposed to MMS alone. The PSM enrichment ratio was increased only for the *PARP1^WT^* but not for the *PARP1^del.p.119K120S^* mutant (2.5 *vs*. 1.1 respectively), indicating efficient trapping. As expected, the profiles of PARP1-interacting proteins in PARP1^WT^-eGFP expressing cells and PARP1^del.p.119K120S^-eGFP expressing cells displayed a number of similarities as well as differences. For example, the trapping-defective PARP1^del.p.119K120S^-eGFP mutant appeared to interact with cohesion complex subunits (SMC1A, SMC3, PDS5A) and other chromatin-associated proteins such as CBX5 (Figure 1F), suggesting that some interaction between mutant PARP1 and chromatin did exist, despite the inability of this PARP1 mutant to be efficiently trapped by PARP inhibitor. Similarly, PARP1^WT^-eGFP appeared to interact with the chromatin-associated proteins (CBX5, BRD3, CHD4), but when compared to the PARP1^del.p.119K120S^-eGFP mutant showed a relative enrichment for interactions associated with the Small Ubiquitin Modifier Proteins, SUMO1 and SUMO2 (Figure 1E). Furthermore, when we compared PARP1^WT^-eGFP interactomes in trapping *vs.* non-trapping conditions (i.e. presence/absence of MMS/PARPi), the SUMO2 PSM ratio increased, suggesting this interaction might be increased when PARP1 is trapped.

As an orthogonal MS approach, we employed Apex2-mediated proximity labelling. Apex2 peroxidase generates free radicals which in the presence of biotin-phenol (BP), biotinylate proteins within a ∼20 nm radius; biotinylated proteins are then purified via Streptavidin-binding. Western blotting confirmed biotinylation of PARP1^WT^-APEX2-eGFP in the presence, but not absence of biotin-phenol, indicating effective labelling (Supplementary Figure 1C). The amount of labelled PARP1 was further increased when PARP1 labelling was conducted under trapping conditions (MMS + talazoparib) (Supplementary Figure 1C). To identify proteins associated with trapped PARP1, we performed Apex2 labelling in cells expressing PARP1^WT^-Apex2-eGFP. As a negative control for the labelling and purification we used PARP1^WT^-eGFP-expressing cells, which because of the absence of Apex2, were unable to perform the biotinylation reaction. Biotinylated proteins were purified under stringent conditions and analysed by mass spectrometry. Non-specific, background protein interactions with beads were removed by filtering the list of PARP1^WT^-Apex2-eGFP-interacting proteins against the list of proteins identified in PARP1^WT^-eGFP expressing cells (detailed analysis description in the Methods). As a result, we identified a higher number of proteins, 360, that associated with PARP1 than for RIME (either in the presence or absence of PARPi, Supplementary Table 3). A STRING network analysis, using a high stringency cut off (0.7) representing the trapped PARP1 interactome network (Supplementary Figure 1D), was enriched in proteins associated with one of the main DNA repair processes PARP1 is involved in, Base Excision Repair (BER), (e.g. PARP1 itself, PCNA, HMGB1, LIG3 and POLE, *p*-value<0.01, Supplementary Figure 1D, E), giving us high confidence in the analysis. Gene Ontology enrichment analysis also identified an enrichment in proteins involved in the spliceosome and ribosome biogenesis (Supplementary Table 4). We also identified a number of well-characterised PARylation targets (e.g. PCNA, NCL, FUS, ILF3^14, 15^) strengthening the notion that we identified *bona fide* PARP1-proximal proteins. Of note, “protein processing in ER” (*p*-value<10^-3^) and “proteasome” (*p*-value<0.01) appeared enriched in the gene set ontology analysis, observations we focus upon later in this manuscript. The MS score and PSM scores showed a positive correlation and identified that among the most abundant proteins were PARP1, p97/VCP, UBA1, TOP2A among others (Figure 1G, H). Proteins that showed high enrichment ratios in PARP trapping conditions e.g., USP7, were generally identified with a low MS score pointing to a low abundance. We prioritised the high MS score over the high PSM ratio in our further considerations as it would represent higher stoichiometric interactions at DNA damage sites.

When we compared the list of proteins identified by RIME with those identified by Apex2 proximity labelling, three proteins were identified by both methods that appeared to have enriched PARP1 interactions under PARP1 trapping conditions: PARP1, p97/VCP and S100A4. The S100A4 protein was identified with a low number of peptides, suggesting a lower stoichiometry interaction, and thus we disregarded it in further analysis. The ATPase p97 attracted our attention as it is a central component of a ubiquitin-controlled process that uses p97’s ATP-dependent unfoldase activity to extract proteins from chromatin prior to their proteasomal degradation or recycling^9, 10, 16^. Furthermore, p97, working with cofactors that often contain ubiquitin binding domains (UBDs), recognises client proteins via ubiquitylation events, mostly those involving lysine-48 (K48) and lysine-6 (K6)^17, 18^ ubiquitylation.

### Trapped PARP1 is sequentially SUMOylated and ubiquitylated

Our RIME analysis suggested that trapped PARP1 might be associated with SUMO1 and SUMO2, whereas our combined RIME/proximity labelling analysis identified p97 as being associated with trapped PARP1. This raised the hypothesis that: (i) trapped PARP1 might be modified by SUMOylation (and possibly consequent SUMO-dependent ubiquitylation); and (ii) these post-translational modifications of trapped PARP1 might, by recruiting ubiquitin-dependent p97, be involved in the processing of trapped PARP1. For example, when we assessed the presence of PARP1 in chromatin-bound and nuclear-soluble fractions from cells cultured MMS + PARPi, we noted the presence of multiple, high molecular weight forms of PARP1 in the chromatin-bound fraction that could conceivably represent SUMOylated and/or ubiquitylated PARP1, which were not present in the nuclear-soluble fraction (Figure 2A).

**Figure 2.**
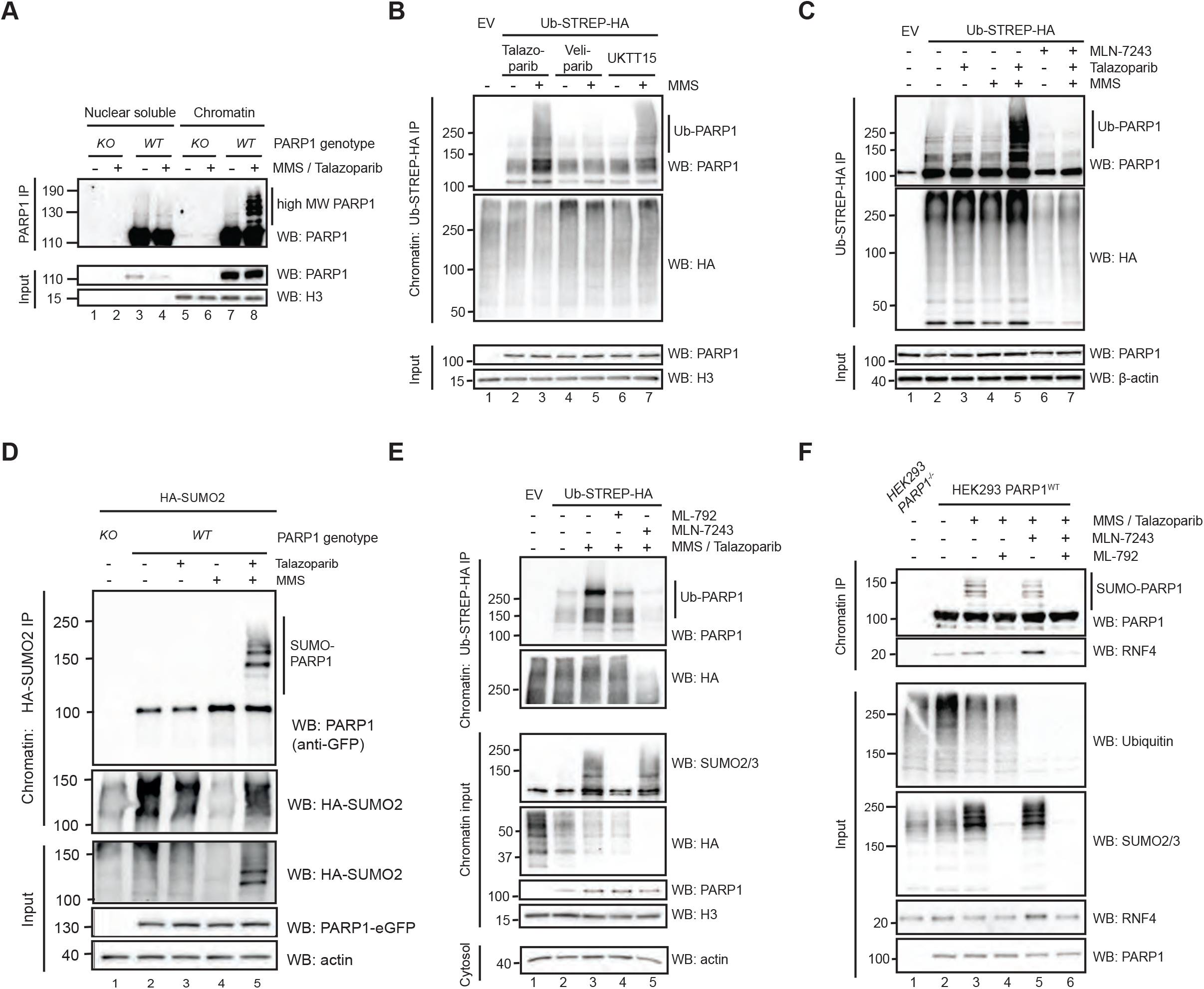
Trapped PARP1 is SUMOylated and ubiquitylated. **A**. Western blot illustrating PARP1 trapping conditions elicit high MW forms of PARP1 in the chromatin fraction. *PARP1-/-* (KO) or *PARP1* wild-type (WT) HEK293 cells were exposed to MMS/talazoparib to stimulate PARP1 trapping, after which cells were fractionated into nuclear-soluble and chromatin-bound fractions. Fractions and whole cell lysates (input) were subjected to PARP1 immunoprecipitation and western blotting for either PARP1 or Histone H3. High MW forms of PARP1 are more prevalent in the chromatin fraction after MMS/talazoparib exposure (lane 8), compared to the nuclear soluble fraction (lane 3,4) or the absence of MMS/talazoparib (lane 3,7). **B.** PARP1 trapping leads to PARP1 ubiquitylation. HEK293 cells were transfected with Ub-STREP-HA-expressing construct and exposed to combinations of 0.01 % MMS, 100 nM talazoparib, 10 μM Veliparib or 10 μM UKTT15 (a veliparib derivative that induces PARP1 trapping). Chromatin fractions were prepared in denaturing conditions to remove protein/protein interactions and immunoprecipitated with streptactin beads to isolate ubiquitylated proteins. Western blotting of Ub-immunoprecipitates indicated that PARP1 is directly ubiquitylated under trapping conditions. The presence of high MW/Ub forms of PARP1 was increased in the presence of the trapping agent UKTT15 (lane 7), while this was not the case with the non-trapping PARPi veliparib (lane 5). **C.** PARP1 trapping conditions result in ubiquitylated PARP1. As in (B), HEK293 cells were transfected with Ub-STREP-HA-expressing construct and exposed to combinations of MMS, talazoparib or 5 μM MLN-7243, a potent ubiquitin E1 ligase inhibitor, as shown. The presence of high MW/Ub forms of PARP1 were reduced by MLN-7243 exposure (lane 7 vs lane 5). Input controls for these experiments are shown in Supplementary Figure 2A. **D.** Trapped PARP1 is SUMOylated. CAL51 PARP1^WT^-eGFP-expressing cells were transfected with HA-SUMO2 expressing construct and subsequently they were treated with 0.01 % MMS or 100 nM talazoparib, either in isolation or in combination. The cells were fractionated and HA-SUMO2-modified proteins were purified from the chromatin fraction. Western blotting for PARP1 showed that high MW isoforms of PARP1 are formed predominantly under trapping conditions. High exposure of blots of PARP1 SUMOylation are shown in Supplementary Figure 2B. **E.** SUMOylation and ubiquitylation inhibitors prevent trapped PARP1 modification. Similarly, to (C), the ubiquitylated pool of proteins was immunoprecipitated from the chromatin fraction of MLN-7243 (5 μM) or ML-792 (1 μM) exposed cells and the presence of high MW PARP1 isoforms identified by immunoblotting. **F**. PARP1 is modified and interacts with RNF4 in a SUMO-dependent manner. HEK293 WT or *PARP1^-/-^* cells were exposed to trapping conditions either in the presence of ubiquitylation (5 μM MLN-7243) or SUMOylation (1 μM ML-792) inhibitors and endogenous PARP1 was immunoprecipitated from the chromatin fraction via PARP1-Trap nanobody. Western blotting for PARP1 revealed that the presence of the modified PARP1 isoforms was abrogated by ML-792 exposure (compare lanes 3 and 4), but not by MLN-7243 exposure (lane 3 vs. 5). This suggests these specific isoforms likely represent SUMOylated PARP1. Abrogating SUMOylation prevented the association between PARP1 and RNF4 (compare lanes 3 and 4), whereas inhibiting ubiquitinating stabilised the interaction.

To extend these observations, we assessed the presence of ubiquitylated PARP1 in cells exposed to PARPi with different trapping properties. In these experiments, we used either the potent PARP1 trapper, talazoparib, or a clinical PARPi that effectively inhibits PARP1 catalytic activity but which has minimal trapping properties, veliparib, or a recently described structural derivative of veliparib, UKTT15, that is able to elicit PARP1 trapping^4^. Cells were exposed to MMS+PARPi, after which the ubiquitylated pool of proteins was isolated from the chromatin fraction via HA-Streptavidin-ubiquitin isolation. Both talazoparib and UKTT15 exposed cells exhibited high molecular weight isoforms of PARP1 in the ubiquitylated pool/chromatin fraction, when compared to cells exposed to veliparib (Figure 2B). This suggested that PARP1 ubiquitylation might be enhanced by PARP1 trapping. To confirm this, we repeated the HA-Streptavidin-ubiquitin pulldown experiment in the presence of the E1 ubiquitin activating enzyme inhibitor, MLN-7243 (TAK243). Western blotting with an anti-PARP1 antibody revealed that trapping conditions led to the formation of high molecular weight PARP1 isoforms; these were almost completely abolished when cells were exposed to MLN-7243, suggesting that these high molecular weight isoforms could represent ubiquitylated PARP1 (Figure 2C, Supplementary Figure 2A). The poly-ubiquitylation of PARP1 was also observed in reciprocal denaturing IP experiments, where PARP1 was immunoprecipitated from HEK293 cells transfected with a FLAG-PARP1 cDNA-expression construct (Supplementary Figure 2B). We also identified poly-ubiquitin chains on PARP1 that were linked by K48 linkage (Supplementary Figure 2C).

The presence of SUMO1 and SUMO2 in our trapped PARP1 interactome, suggested PARP1 may also be modified by SUMOylation. We therefore reasoned that the high molecular PARP1 modifications could also represent SUMOylated, in addition to ubiquitylated PARP1. To test this hypothesis, we expressed HA epitope-tagged SUMO2 and isolated the SUMOylated pool of proteins under denaturing conditions from the chromatin fraction and found that trapped PARP1 was heavily modified by SUMOylation (Figure 2D). We also found that when cells were exposed to MMS alone (to induce DNA damage and activate PARP1) in the absence of PARPi, there was a depletion in the total pool of SUMO2 and a minimal level of PARP1 SUMOylation, as previously observed^19^ (Figure 2D, Supplementary Figure 2D). However, this was not to the same extent as seen under PARP1 trapping conditions. Taken together with our prior observations, this suggested that PARP1 SUMOylation might be important when PARP1 is trapped. Interestingly, incubating cells grown in PARP1 trapping conditions in the presence of a SUMOylation inhibitor (ML-792) decreased high MW forms of ubiquitylated PARP1 (Figure 2E), suggesting PARP1 ubiquitination upon trapping could require prior PARP1 SUMOylation. Conversely, the inhibition of ubiquitylation had no effect on PARP1 SUMOylation (Figure 2F). These observations suggested that the SUMOylation of trapped PARP1 is required for its ubiquitination, but that the ubiquitination of trapped PARP1 is not a pre-requisite for PARP1 SUMOylation.

### Trapped PARP1 is sequentially SUMOylated by PIAS4 and ubiquitylated by RNF4

The SUMOylation followed by ubiquitylation of trapped PARP1 suggested the concert action of a SUMO E3 ligase and a SUMO-targeted ubiquitin ligase (STUbL). In the first instance, we assessed the roles of PIAS4 and RNF4 as candidates, as PIAS4 has been previously implicated as a SUMO E3 ligase for PARP1 in its non-trapped state^20^ and RNF4, a STUbL, has previously been implicated in modulating PARP1’s transcriptional activity^21^ and in repairing topoisomerase cleavage complexes, which also represent a “trapped” nucleoprotein complex^22^. Indeed, chromatin co-immunoprecipitation of trapped PARP1 showed increased interaction with RNF4. Importantly, this interaction was reduced upon SUMOylation inhibition and stabilised upon ubiquitination inhibition (Figure 2F), indicative of a ligase-substrate interaction.

To delineate the relationship between SUMOylation and ubiquitylation of trapped PARP1 and a possible role for PIAS4 in this process, we used HCT116 *PIAS4^−/−^* and MCF7 *RNF4^−/−^* cell lines^22^. Both cell lines were transfected with a FLAG-PARP1-expressing cDNA construct after which PARP1 was immunoprecipitated from the chromatin fraction of cells grown in trapping conditions. Western blotting with an anti-SUMO2/3 antibody revealed that PIAS4 activity is necessary for efficient SUMOylation and ubiquitylation of trapped PARP1 (Figure 3A, Supplementary Figure 3A and B). Re-expressing PIAS4 ^WT^ in HCT116 *PIAS4^−/−^* cells reversed these effects, but this was not achieved when we expressed a DNA-binding SAP domain deleted or the catalytically inactive C342A PIAS4 mutant^23^ (Figure 3B and C). Interestingly, in *RNF4^−/−^* cells, while PARP1 ubiquitylation was decreased, confirming that RNF4 activity is responsible for this modification, SUMOylation was increased (Figure 3D, Supplementary Figure 3D and E). Re-expressing RNF4^WT^ in MCF7 *RNF4^-/-^* cells reversed these effects, but this was not the case for SIM (SUMO-interacting motifs)-deleted^24^ or catalytically inactive H156A mutant (Figure 3E and F). Using gene silencing, we again found that RNF4 activity loss reduced ubiquitylation of trapped PARP1 (Supplementary Figure 3G), establishing RNF4 as a STUbL E3 ligase for trapped PARP1. It is important to point out, that in both *PIAS4^-/-^* and *RNF4^-/-^* cells, there existed some amount of residual SUMOylation and ubiquitylation, suggesting that other E3 SUMO and ubiquitin ligases might have a role here.

**Figure 3.**
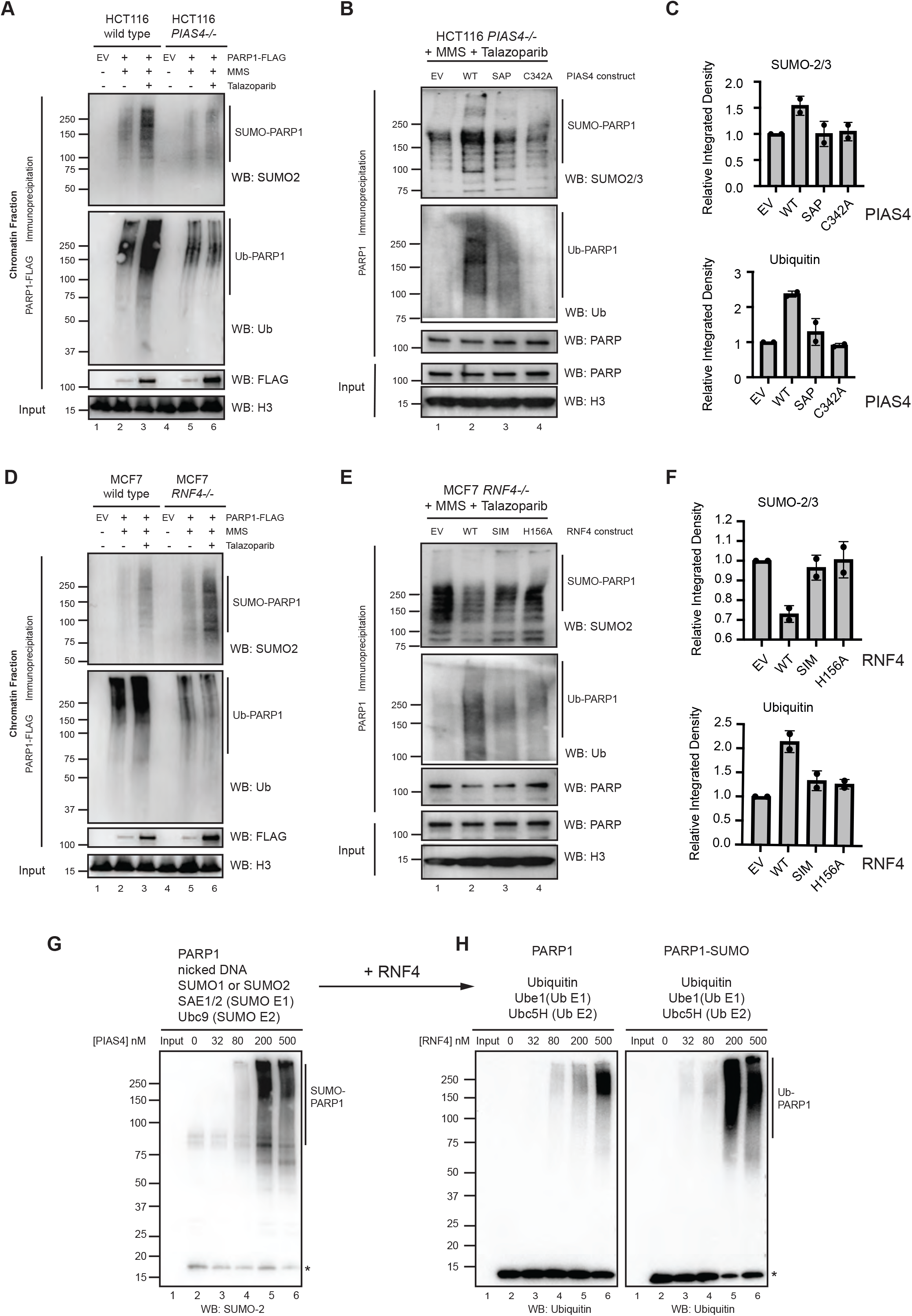
Trapped PARP1 is modified in a PIAS4- and RNF4-dependent manner. **A**. PARP1 is SUMOylated in a PIAS4-dependent manner *in vivo*. HCT116 wild-type or *PIAS4*^−/−^ cells were transfected with FLAG-PARP1 expressing plasmid and subsequently cultured in PARP1-trapping conditions. FLAG immunoprecipitation was carried out from the chromatin-bound fraction (i.e. trapped PARP1 fraction) and the presence of SUMOylation and ubiquitylation was detected via western blotting. In *PIAS4*^−/−^ cells, the amount of SUMO1 (Supplementary Figure 3A) and SUMO2 was significantly reduced. Similarly, ubiquitylation was also reduced. Quantification of the blots is presented in Supplementary Figure 3B. **B**. HCT116 *PIAS4*^−/−^ cells were transfected with indicated PIAS4-expressing plasmids (EV: empty vector, WT: wild type, *SAP*: SAP domain deleted, C342A catalytic dead) for 48 hours, followed by 30 min talazoparib (10 µM) treatment in the presence of 0.01 % MMS and whole cellular immunoprecipitation using PARP1 antibody. Immunoprecipitate and input were subjected to Western blotting and detected using indicated antibodies. **C**. A quantification of the abundance of SUMO2/3 (top) and ubiquitin (bottom) modified PARP1 isoforms in the cells transfected as in (B). Two biological replicates are shown. **D**. Similar to (A), trapped PARP1 was purified from MCF7 wildtype or *RNF4*-/-cells. In *RNF4*-/-cells the amount of ubiquitylation was reduced, whilst the amount of SUMO1 (Supplementary Figure 3D) and SUMO2 was increased. Quantification of the blots is presented in Supplementary Figure 3E. **E**. MCF7 *RNF4*^−/−^ cells were transfected with indicated RNF4-expressing plasmids (EV: empty vector, WT: wild type, *SIM*: SUMO-interacting motif deleted, H156A catalytic dead) for 48 hours, followed by 30 min talazoparib (10 µM) treatment in the presence of 0.01 % MMS and whole cellular immunoprecipitation using PARP1 antibody. Immunoprecipitate and input were subjected to Western blotting and detected using indicated antibodies **F**. A quantification of the abundance of SUMO2/3 (top) and ubiquitin (bottom) modified PARP1 isoforms in the cells transfected as in (E). Two biological replicates are shown. **G**. PIAS4 mediates PARP1 SUMOylation *in vitro*. Recombinant PARP1 was incubated in the presence of nicked DNA, SUMO1 or SUMO2, SAE1/2 (SUMO E1), Ubc9 (SUMO E2) and an increasing concentration of PIAS4. PIAS4 led to a concentration-dependent increase of SUMOylated PARP1, as detected by anti-SUMO2 or anti-SUMO1 (Supplementary Figure 3H) immunoblotting. Pool of free SUMO2 is indicated by*. **H**. RNF4 mediates PARP1 ubiquitylation in a SUMO-dependent manner *in vitro*. Similar to (H), PARP1 SUMOylation reactions were supplemented with ubiquitin, Ube1 (E1), Ubc5H (E2) and an increasing concentration of RNF4. The addition of RNF4 led to a concentration-dependent ubiquitylation as detected by anti-ubiquitin antibody. PIAS4 SUMOylated PARP1 was a better substrate for ubiquitylation. Pool of free ubiquitin is indicated by*.

Finally, we tested the interdependency of PARP1 SUMOylation and ubiquitylation events using *in vitro* SUMOylation and ubiquitylation reactions. Incubating PARP1 in the presence of nicked DNA, SUMO1 or SUMO2, SAE1 (SUMO E1), Ubc9 (SUMO E2) and increasing concentration of PIAS4 led to a concentration-dependent formation of SUMO-modified PARP1 (Figure 3G and Supplementary Figure 3H). Further addition to the reactions with ubiquitin, UBE1 (Ub E1), Ubc5H (Ub E2) and an increasing concentration of RNF4, led to efficient PARP1 ubiquitylation (Figure 3H). In contrast, RNF4 displayed much lower ubiquitylating activity towards PARP1 in the absence of SUMOylation (Figure 3H). Collectively, these data suggested a stepwise process, where upon trapping, PARP1 is initially SUMOylated by PIAS4, followed by STUBL RNF4 driven ubiquitylation.

### p97 interacts with modified trapped PARP1

Although the above experiments suggested a link between the trapping of PARP1 by PARPi, PARP1 SUMOylation and ubiquitylation, the functional significance of these events remained to be determined. Our mass-spectrometry analysis also suggested that under PARP1 trapping conditions, PARP1 might interact with p97 ATPase, a central component of the ubiquitin system involved in chromatin-associated degradation. We therefore hypothesised that the SUMOylation and ubiquitylation of trapped PARP1 might be essential for the recruitment of p97 ATPase to process trapped PARP1, thus removing it from DNA breaks.

We first confirmed the interaction between p97 and PARP1 using both Proximity Ligation Assays (PLA) (Figure 4A) and co-immunoprecipitation experiments (Supplementary Figure 4A). Both experimental approaches, verified that this interaction was enhanced in a trapping-dependant manner in PARP1 wild-type cells but not in those with a DNA binding-deficient PARP1 mutation (Figure 4B, C). Exposing cells to UKTT15, but not veliparib, also led to an increase in the PARP1-p97 interaction, validating the importance of trapping for this interaction, as opposed to catalytic inhibition of PARP1 with minimal trapping, as achieved with veliparib (Supplementary Figure 4B).

**Figure 4.**
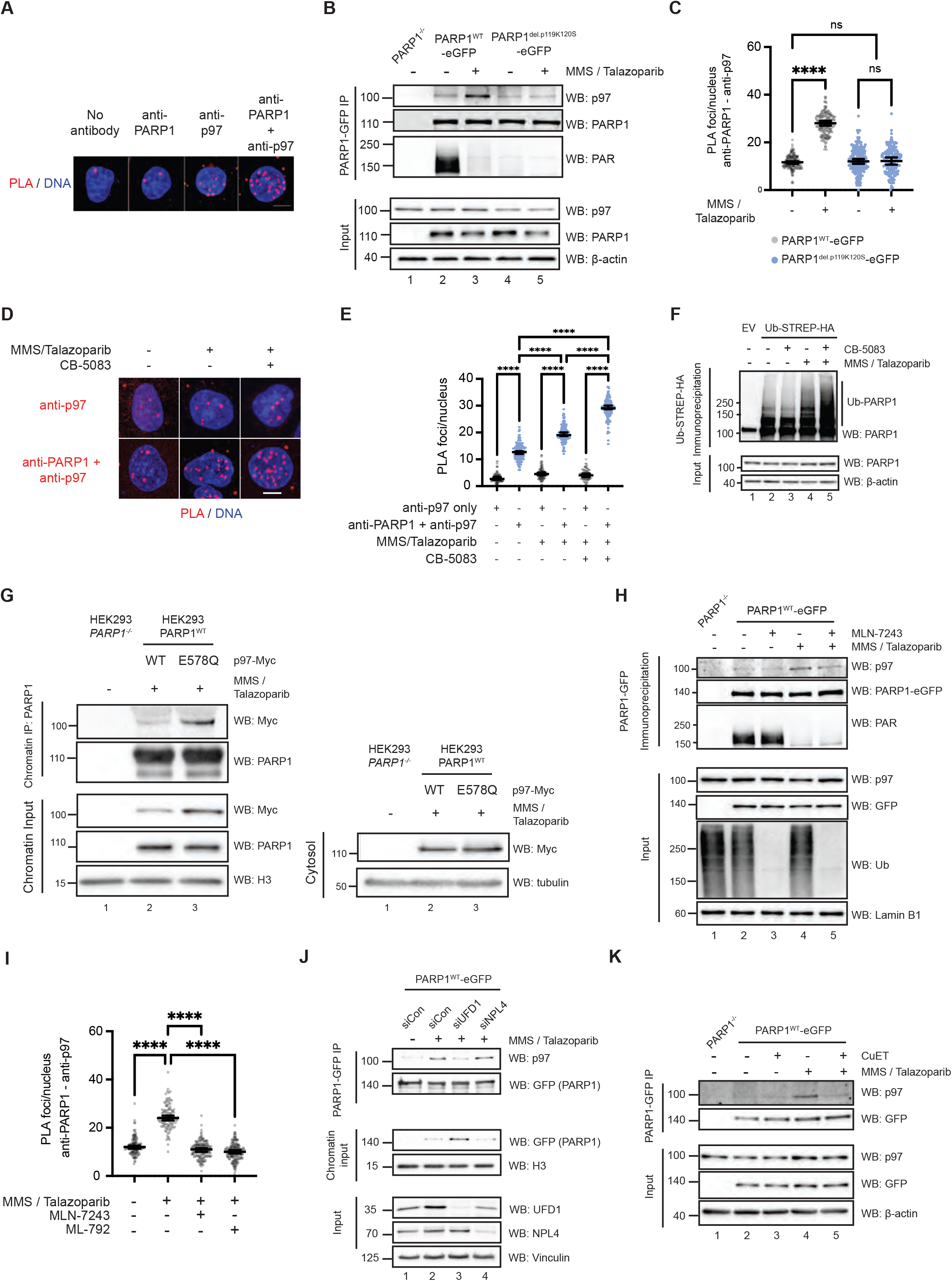
PARP1 interacts with p97 in a trapping-dependent manner. **A.** Confocal microscopy images from a Proximity Ligation Assay (PLA) experiment monitoring co-localisation of endogenous PARP1 and p97 in CAL51 cells. Nuclei (blue) were visualised by DAPI staining and PARP1/p97 co-localisation by PLA (red). Nuclear PLA foci were detected when both PARP1 and p97 antibodies were used. PLA foci are quantified in panel (C). Representative images are shown, scale bar = 5 μm. **B**. PARP1-p97 interaction is increased upon DNA damage. PARP1^WT^-eGFP or PARP1^del.pK119S120^-eGFP expressing CAL51 cells were exposed to trapping conditions. Cells were then lysed and chromatin digested by benzonase. PARP1-GFP was immunoprecipitated from lysates under native conditions (150 mM NaCl, 50 mM Tris-HCl) and then western blotted as shown. The p97/PARP1 interaction in PARP1^WT^-eGFP expressing cells was enhanced under trapping conditions (lane 2 *vs.* 3); this was not observed in cells expressing the PARP1^del.pK119S120^ mutant (lanes 4,5). **C.** PARP1 and p97 interact in a trapping-dependent manner. PARP1-p97 PLA analysis is shown from CAL51 cells expressing either PARP1^WT^-eGFP or PARP1^del.pK119S120^-eGFP. Compared to non-trapping conditions, MMS/PARPi exposure increased the PARP1-p97 PLA signal in cells expressing PARP1^WT^-eGFP but not in those expressing PARP1^del.pK119S120^-eGFP. Quantification of the number of PLA foci/nucleus in n>200 cells from three independent experiments; black bars show the geometric mean ± 95CI, *p*-values were calculated with one-way ANOVA, **** - *p* < 0.0001. **D.** Confocal microscopy images from a PARP1/p97 PLA experiment where CAL51 cells were exposed to combinations of 0.01 % MMS and 100 nM talazoparib (to induce trapping). PLA was conducted either with p97 antibody only (top row) or p97+PARP1 antibody (bottom row). P97 inhibitor (10 μM CB-5083) was added to stabilise the interaction between p97 and its substrate. Representative images are shown, scale bar = 5 μm. **E.** Quantification of PLA foci/nucleus from experiments in (D). The combination of MMS/talazoparib and CB-5083 enhanced the nuclear PARP1/p97 PLA signal. Quantification of the number of PLA foci/nucleus in n>200 cells from three independent experiments; black bars show the geometric mean ± 95CI, *p*-values were calculated with one-way ANOVA, **** - *p* < 0.0001. **F**. p97 inhibition increases the presence of ubiquitylated PARP1. Ubiquitin-STREP-HA-expressing HEK293 cells were cultured in PARP1 trapping conditions in the presence or absence of 10 µM CB-5083; the pool of ubiquitylated proteins was then immunoprecipitated under denaturing conditions and the presence of PARP1 in the immunoprecipitates detected by immunoblotting. Input controls for these experiments are shown in Supplementary Figure 4C. **G.** HEK293 cells expressing doxycycline-inducible p97-Myc constructs (wither wild-type or dominant negative E578Q mutant) were transfected with a FLAG-PARP1-expressing constructs. After induction with 1 µg/ml doxycycline for 16 h and subsequently treated with trapping conditions, the cells were fractionated and PARP1 was immunoprecipitated with PARP-Trap beads from the chromatin-bound fraction. The expression of the dominant negative E578Q mutant leads to stronger interaction with PARP1 in the chromatin. **H.** Ubiquitin is required for the PARP1/p97 interaction in trapping conditions. Western blots of PARP1 co-immunoprecipitates from CAL51 PARP1^WT^-eGFP-expressing cells are shown. Exposure to trapping conditions increases the PARP1/p97 interaction (lane 4), an effect reversed by MLN-7243 (5 μM). PARP1-GFP was immunoprecipitated under native conditions using GFP-Trap beads and the presence of p97 was detected by western blotting. **I.** PARP1/p97 co-localisation is reduced by ubiquitylation (5 μM MLN-7243) or SUMOylation (1 μM ML-792) inhibitors. Quantification of PARP1/p97 PLA foci/nucleus from CAL51 cell cultured in MMS/talazoparib plus the corresponding inhibitors. Either MLN-7243 or ML-792 reduces the number of PLA foci/nucleus. Quantification of the number of PLA foci/nucleus in n>200 cells from three independent experiments; black bars show the geometric mean ± 95CI, *p*-values were calculated with one-way ANOVA, **** - *p* < 0.0001. **J**. The p97 adapter UFD1 mediates the interaction with trapped PARP1. PARP1-p97 interaction was investigated by Co-IP from chromatin-bound (trapped) PARP1. RNAi-mediated UFD1 depletion strongly reduced the amount of p97 recruited to the trapped PARP1 complex and caused increased total levels of trapped PARP1 in the chromatin fraction. This was not the case in Npl4-depleted cells. **K.** The PARP1/p97 interaction is disrupted by the p97 sequestration agent, CuET. Similar to Co-IPs shown above, western blot of PARP1 immunoprecipitates from CAL51 PARP1^−/−^ cells stably expressing PARP1^WT^-eGFP are shown. Exposure to trapping conditions increases the PARP1/p97 interaction (lane 4), an effect reversed by CuET exposure (1 μM, lane 5).

The inhibition of p97 ATPase activity by the inhibitor CB-5083, which induces a p97 substrate trapping effect^25^, caused an increase in the interaction between p97 and PARP1 (Figure 4D, E), suggesting that PARP1 could be a p97 substrate. Blocking p97 catalytic activity leads to accumulation of ubiquitylated isoforms of its substrates ^26, 27^, which was also the case for trapped PARP1 (Figure 4F). Using reciprocal immunoprecipitation under denaturing conditions, we again observed increased PARP1 ubiquitylation upon trapping and p97 inhibition by CB-5083 (Figure 4F, Supplementary Figure 2C, D). Furthermore, we reproduced the substrate trapping effect of p97 inhibition by expressing a ATPase deficient p97 (E578Q) mutant, which acts in a dominant negative manner^16, 28, 29^ in HEK293 cells. The p97-E578Q mutant showed a stronger interaction with chromatin-associated PARP1 under trapping conditions, consistent with trapped PARP1 being a p97 substrate (Figure 4G). Furthermore, by expressing the p97-E578Q mutant in the PARP1-reconstituted CAL51 cells, we demonstrated that the p97-PARP1 interaction was trapping-dependent as it was present in cells, expressing PARP1^WT^, but not in cells expressing the trapping-deficient PARP1^del.pK119S120^ mutant (Supplementary Figure 4D).

Ubiquitylation is a mediator of p97 interactions. Indeed, when cells were exposed to ubiquitylation (MLN-7243) or SUMOylation (ML-792) inhibitors (which decreased trapped PARP1 ubiquitylation), the interaction between PARP1 and p97 was also reduced, as measured by co-precipitation (Figure 4H) or PLA (Figure 4I). p97 recognises and processes its ubiquitylated substrates using a series of cofactors, including the NPL4-UFD1 complex, which mostly serves as a ubiquitin binding receptor due to ubiquitin-binding domains (UBDs) in both NPL4 and UFD1^30, 31^. When UFD1 was depleted by RNA interference, the interaction between trapped PARP1 and p97 was reduced (Figure 4J). Surprisingly, this was not the case when NPL4 was depleted (Figure 4J). Furthermore, depletion of UFD1 lead to a strong accumulation of trapped PARP1 in chromatin (Figure 4J and Supplementary Figure 5D). Our observations appear consistent with previous work suggesting that the NPL4 and UFD1 can recognise substrates independently of each other^7, 32,9^. We also evaluated the effect of CuET, a metabolite of the alcohol-abuse drug disulfiram, which acts to specifically segregate the UFD1-NPL4-p97 complex from chromatin into agglomerates^33^; in this way, CuET has a distinct mechanism of action compared to CB-5083, which inhibits p97 ATPase activity on the entire cellular p97 pool as opposed to chromatin associated p97-UFD1-NPL4 complex. We found that the PARP1-p97 interaction was almost completely abrogated by CuET exposure (Figure 4K).

Taken together, these observations suggested that the p97 system, including p97 ATPase activity and the ubiquitin binding cofactor UFD1 (p97-UFD1), recognises and physically interacts with trapped PARP1.

### Trapped PARP1 is modulated by p97 activity

Given that p97 interacts with PARP1, we hypothesised that p97 removes trapped PARP1 from chromatin. To assess this, we used a “trap-chase” experimental approach to monitor the kinetics of how trapped PARP1 is resolved after cells are exposed to PARPi (Figure 5A). Cells were exposed to MMS/PARPi to induce trapping (the “trap”) for a defined period and subsequently cultured in either fresh media with no MMS/PARPi or in media containing different combinations of PARPi and p97 complex inhibitors (the “chase”). At various time points during the chase, the amount of trapped PARP1 was evaluated either by chromatin fractionation or by a recently described method that estimates the amount of DNA damage-associated PARP1 by measuring the proximity of PARP1 with phosphorylated H2AX (γH2AX) by PLA^34^.

**Figure 5.**
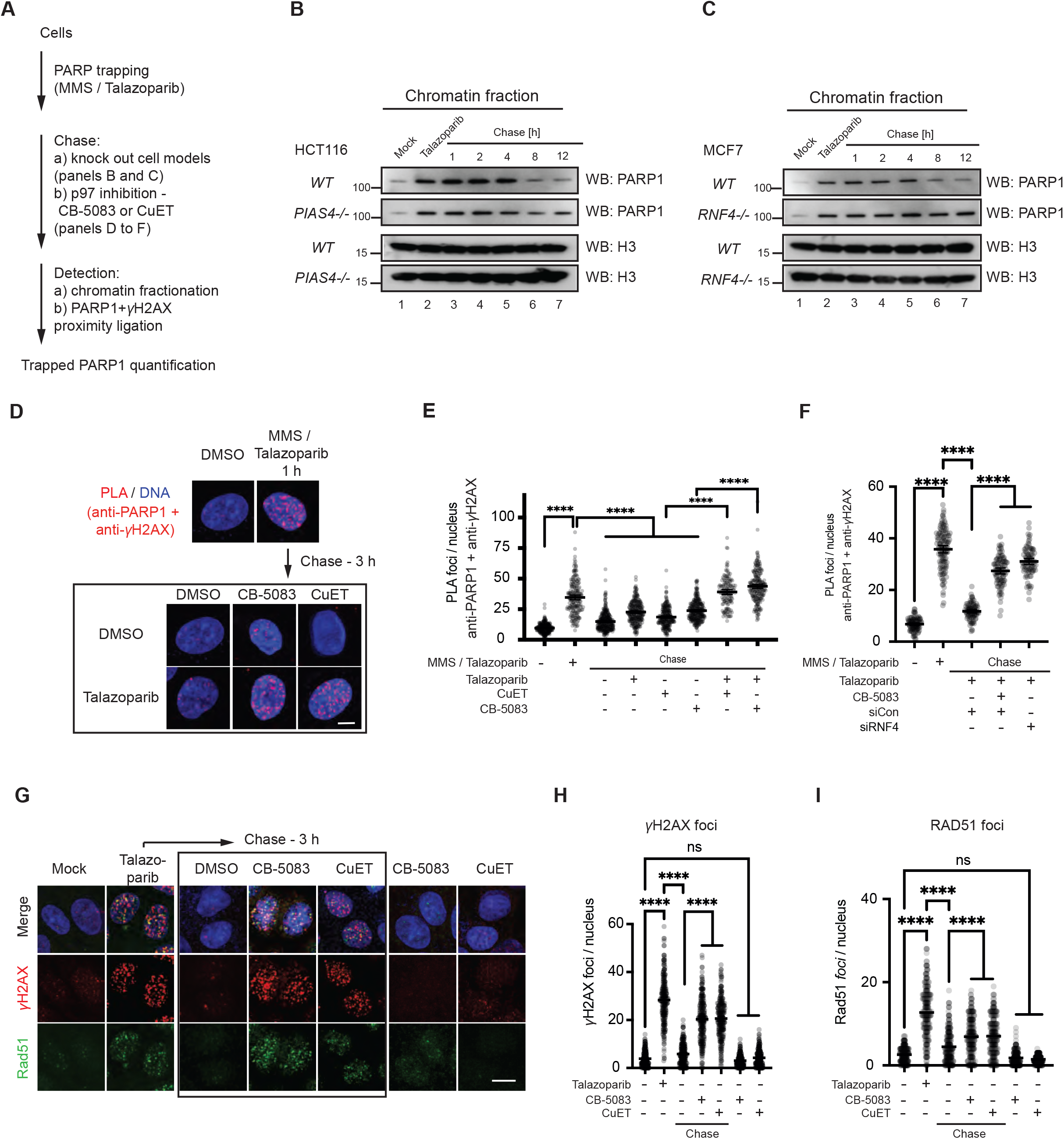
PARP trapping is modulated by the PIAS4-RNF4-P97/VCP axis. **A.** Schematic of the trap-chase experiment is shown. Cells were cultured in trapping conditions (0.01% w/v MMS + 100 nM talazoparib) (trap), washed and then cultured in talazoparib containing media (chase). The chase was done either in KO cell models or in combination with media containing p97 inhibitors (CB-5083 or CuET), for a subsequent period. Prior to- and then after both trap and chase, the extent of trapped PARP1 was detected by both chromatin fractionation or PARP1/γH2AX PLA. **B.** Trapped PARP1 is processed in a PIAS4-dependent manner. A trap-chase experiment in HCT116 wild-type or *PIAS4^−/−^* cells. After trapping by 100 nM talazoparib + 0.01% MMS, cells were chased in a talazoparib-containing media and collected at indicated time points, followed by chromatin fractionation and Western blotting. The amount of chromatin-bound PARP1 decreased over the chase period. In the *PIAS4^−/−^* cells, this process was delayed in the later time points. **C**. Trapped PARP1 is processed in a RNF4-dependent manner. A trap-chase experiment in MCF7 wild-type or *RNF4^−/−^* cells similar to (B). The amount of chromatin-bound PARP1 decreases over the chase period. In *RNF4^−/−^* cells this process was delayed. **D.** Representative confocal microscopy images from a PARP1/γH2AX PLA trap-chase experiment. Nuclei, stained with DAPI, shown in blue, PARP1/γH2AX PLA foci shown in red. Scale bar = 5 μm. **E.** PARP1/γH2AX PLA foci persist in cells chased in PARPi plus p97 inhibitors. Quantification of PARP1/γH2AX PLA foci from the trap-chase experiment in (D). Trapping conditions (MMS+talazoparib) increase PARP1/γH2AX PLA foci which decline after a chase in talazoparib alone or the absence of additional small molecule inhibitors. PARP1/γH2AX PLA foci persisted when cells were chased in talazoparib in combination with either CB-5083 or CuET. Quantification of the number of PLA foci/nucleus in n>200 cells from three independent experiments; black bars show the geometric mean ± 95CI, *p*-values were calculated with one-way ANOVA, **** - *p* < 0.0001. **F.** PARP1/γH2AX PLA foci persist in cells with RNF4 silencing. CAL51 cells were transfected with siRNA targeting RNF4. 48 hours post RNF4 depletion, a PLA trap-chase experiment (PARP1-γH2AX) was performed. 10 μM CB-5083 in the chase was used as a control for blocking p97 activity. Quantification of the number of PLA foci/nucleus in n>200 cells from three independent experiments; black bars show the geometric mean ± 95CI, *p*-values were calculated with one-way ANOVA, **** - *p* < 0.0001. **G.** PARPi-induced RAD51 and DH2AX foci persist in the presence of p97 inhibitors. Representative confocal microscopy images from a trap-chase experiment (where cells were incubated with talazoparib overnight) are shown, where the effect of PARPi was monitored by immunofluorescent detection of γH2AX and RAD51 foci. In the chase phase the PARPi was washed out and the cells were chased in control or p97 inhibitor-containing media. Representative images for each condition, scale bar = 5 μm. **H** and **I**. Quantification of γH2AX (H) and RAD51 foci (I), from experiment (G). Quantification of the number of foci/nucleus in n>200 cells from three independent experiments; black bars show the geometric mean ± 95CI, *p*-values were calculated with one-way ANOVA, **** - *p* < 0.0001.

Initially, we followed the kinetics of trapped PARP1 in *PIAS4^−/−^* and *RNF4^−/−^* cells that have reduced trapped PARP1 ubiquitylation (Figure 3A and 3D). These cells were exposed to trapping conditions and chased in talazoparib-containing media. *PIAS4^−/−^* (Figure 5B) and *RNF4^−/−^* (Figure 5C) cells showed slower resolution of chromatin bound PARP1, especially at the later time points, consistent with the notion that these SUMO/ubiquitin ligases, promote the resolution of trapped PARP1 complex.

Secondly, we investigated the role of p97 activity. We conducted a trap-chase experiment and included talazoparib and CB-5083 or CuET in the chase phase of the experiment. We monitored the amount of trapped PARP1 either via chromatin fractionation (Supplementary Figure 5A) or PLA (Figure 5D). After exposing cells to MMS/talazoparib, a significant amount of PARP1 was detected in the proximity of γH2AX (Figure 5D), indicating the “trapping” part of the experiment was successful; after removing the trapping agents by washing cells, the amount of trapped PARP1 decreased (e.g. the PARP1/DH2AX PLA signal disappeared). When cells were chased in the presence of single agent PARPi or p97 inhibitor, the PARP1/γH2AX PLA signal also diminished. Conversely, when cells were chased in the presence of both, PARPi (talazoparib) and p97 inhibitor (either CB-5083 or CuET), the amount of trapped PARP1 persisted (Figure 5D, E). Consistent with the notion that RNF4 is an upstream factor involved in the processing of trapped PARP1, we also found that gene silencing of RNF4 led to the persistence of PARP1/γH2AX PLA foci (Figure 5F). Additionally, we assessed the effect on PARP trapping by the expression of a dominant negative RNF4-M136S/R177A mutant (Supplementary Figure 5B), a p97-E578Q mutant (Supplementary Figure 5C) or UFD1 depletion (Supplementary Figure 5D). All three led to higher level of trapped PARP1 in the chromatin fraction under trapping conditions, confirming the importance of these proteins in the processing of trapped PARP1.

In homologous recombination proficient cells trapped PARP1 activates RAD51-mediated DNA repair; this latter event can be monitored by assessing the formation of nuclear RAD51 foci. As expected, we found that a 16 hour exposure of cells to PARPi elicited both γH2AX and RAD51 foci responses (Figure 5G and quantified in Figure 5H and I). When PARPi was removed from culture media by washing, γH2AX and RAD51 foci disappeared after three hours, suggesting resolution of the DNA damage caused by trapped PARP1.

However, when we used p97 inhibitors (CB-5083 or CuET) in this “chase” period, γH2AX and RAD51 foci persisted, indicating that the underlying trapped PARP1-induced damage could not be resolved efficiently. Notably, incubating the cells in the presence of single-agent p97 inhibitors did not induce γH2AX and RAD51 foci, suggesting that the persistence of γH2AX and RAD51 foci in experiments involving PARPi plus p97 inhibitor were indeed caused by PARPi. The effects in foci resolution were also not trivially explained by alterations in the cell cycle as three hours incubation of cells in p97 inhibitor did not lead to significant changes in cell cycle distribution (Supplementary Figure 5E, F).

The above observations suggested that p97 activity is critical for removing trapped PARP1. Recently, Shao et al, by using fluorescence recovery after photobleaching (FRAP), have demonstrated that trapped PARP1 molecules are exchanged at the sites of DNA damage, even in the presence of trapping PARPi^35^. This raised the possibility that p97 is one of the factors that facilitates this exchange. To address this possibility, we conducted FRAP experiments and observed consistent FRAP signal recovery at UV-laser-induced DNA damage sites (Supplementary Figure 5G, H). The addition of p97 inhibitor (CB-5083) led to a modestly slower FRAP (from talazoparib *t_1/2_* = 4.9 ± 1.3 s to talazoparib + CB-5083 *t_1/2_* = 7.8 ± 1.4 s, two-sided *t*-test *p*-value < 0.05), but did not abrogate it altogether. These results suggested that the role that p97 has on PARP1 trapping cannot be entirely recapitulated at UV-laser-induced lesion sites, indicating that there may exist distinct molecular events at different DNA damage sites that lead to formation of PARP1 trapping that are resolved by p97.

### p97 inhibition potentiates PARP inhibitor cytotoxicity

Based on the prolonged PARP1 trapping effects described above, we also hypothesised that p97 inhibition might modulate the cytotoxic effects of PARP inhibitors. Initially, we assessed the effect of two p97 inhibitors (CB-5083 and CuET) on the cytotoxic effect of two trapping PARPi (talazoparib and olaparib). We observed a dose dependent potentiation of the effect of the PARPi by the presence of p97 inhibitor in long-term clonogenic survival experiments (Figure 6A, B and C). This effect was trapping dependent as it was reversed in *PARP1^-/-^* cells (Figure 6A and Supplementary Figure 6A and B), suggesting that it was also not due to other roles that p97 might play in DNA repair. Furthermore, when cells were exposed to alkylating agents (MMS or temozolomide) used in combination with p97 inhibitor, no sensitisation was observed in either PARP1^WT^ or *PARP1^-/-^* cells (Figure 6D, E), implying that other roles p97 might play in alkylation DNA damage repair are unlikely to explain the PARPi sensitisation.

**Figure 6.**
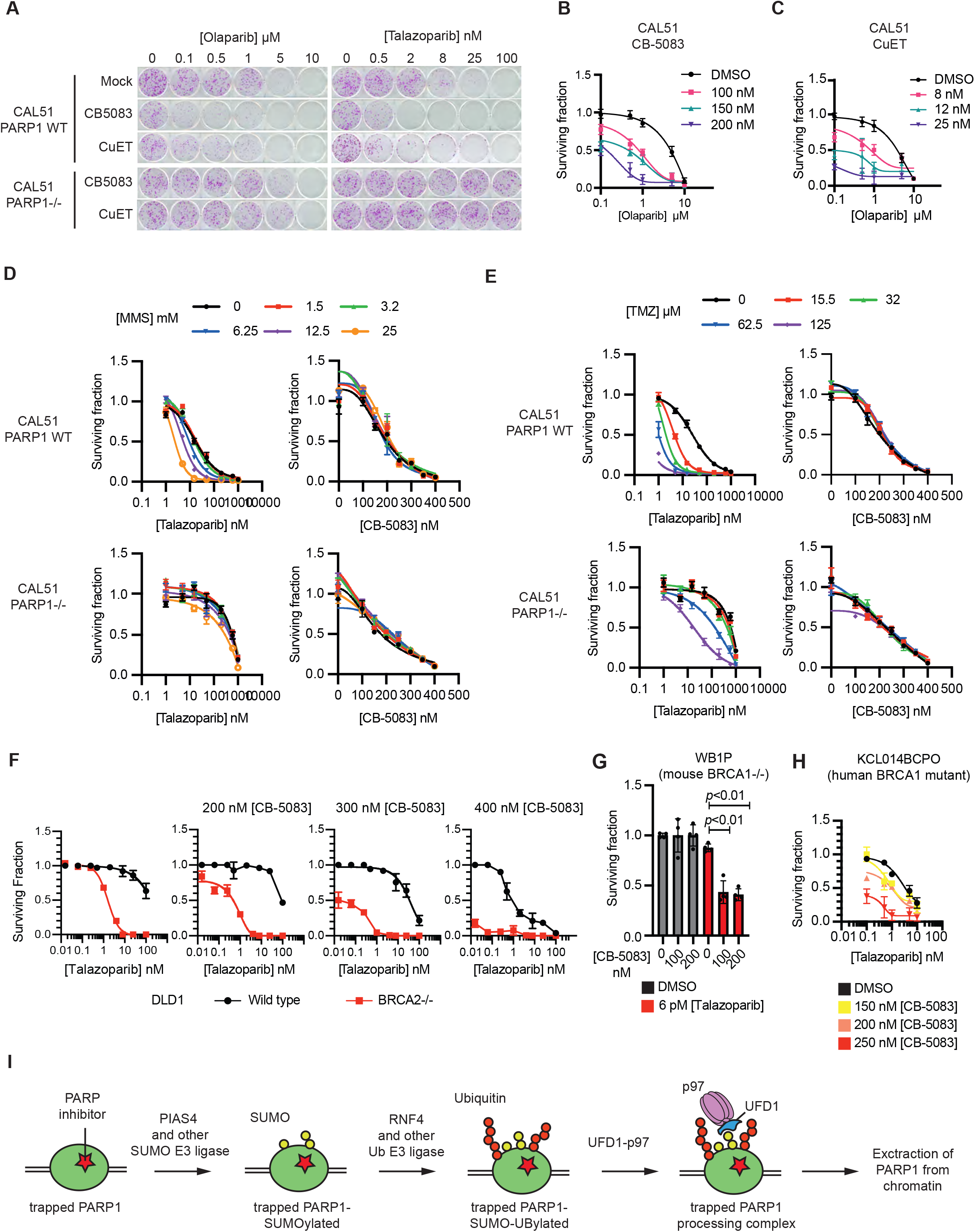
p97 inhibition potentiates the effect of PARP inhibitors. **A.** p97 inhibition potentiates the cytotoxicity of PARP inhibitors. CAL51 cells were treated with increasing PARP inhibitor concentration (Talazoparib or Olaparib) in the presence of increasing p97 inhibitor concentrations (CB-5083 or CuET) for a period of 14 days. Subsequently the colonies were fixed and stained by sulphorhodamine B. The presence of both p97 inhibitors led to the potentiation of the PARPi cytotoxic effect. Shown are images for the 100 nM CB-5083 and 8 nM CuET treatment samples. This effect was reversed in *PARP1^-/-^* CAL51 cells, treated with the same concentrations, showing that it is PARP1-mediated synthetic lethality. Drug response curves are shown in (B), (C) and supplementary figure 6A, B. **B.** and **C.** Drug-response curves for colony formation experiments shown in (A). **D.** and **E**. DNA alkylating agents that are used to induce trapping do not synergise with p97 inhibition in the absence of trapping PARP inhibitor. CAL51 WT or *PARP1^-/-^* cells were treated with increasing concentrations the alkylating agents MMS (D) or temozolomide (TMZ) (E) in combination with either trapping PARP inhibitor or p97 inhibitor. After seven-day exposure, cellular viability was measured by CellTiter Glo. PARPi showed strong sensitizing effect to alkylating agents in PARP1 WT cells. p97 inhibitor did not demonstrate any sensitizing effect in either WT or *PARP1^-/-^* cells, suggesting that alkylating damage alone cannot sensitise to p97 inhibition and that trapped PARP1 is required for this effect. **F**. CB-5083 modulates the synthetic lethal effect of PARPi in *BRCA2^−/−^* cells. Survival curves from clonogenic survival assays in DLD1 *BRCA2^wild-type^* and DLD1 *BRCA2^−/−^* cells. Colony formation images and quantification of the Area Under the Curve (AUC) are shown in Supplementary Figure 6E. **G.** p97 inhibition sensitises mouse cancer organoid cells to PARPi. *Brca1/p53* mutant WB1P breast cancer organoids were grown in the presence of low concentration PARPi (0.006 nM talazoparib) which when used as a single agent had no impact on cell survival. CB-5083 was added to a final concentration of 100 or 200 nM and organoids cultured in a 3D culture for seven days. Viability was evaluated by 3D CellTiter-Glo and quantification of the Surviving Fraction is presented. Brightfield images of organoids are shown in Supplementary Figure 6F. **H**. p97 inhibition sensitises a human *BRCA1* mutant patient-derived breast cancer organoid to PARPi. KCL014BCPO organoids were grown in increasing concentrations of talazoparib and CB-5083 for a period of seven days. Viability was evaluated by 3D CellTiter-Glo and quantification of the Surviving Fraction is presented. Brightfield images of organoids are shown in Supplementary Figure 6G. **I.** A model of the processing of trapped PARP1. PARP1 trapped by the presence of PARPi on DNA is processed in a stepwise manner. It is initially SUMOylated in a PIAS4-dependant manner and subsequently ubiquitylated in an RNF4-dependent manner. p97 is recruited to the ubiquitin chains and binds via UFD1 and the ATPase activity of p97 extracts the modified PARP1 from the chromatin.

Because PARPi are approved for the treatment of cancers that have homologous recombination defects, including those with *BRCA1* or *BRCA2* gene defects, and because trapped PARP1 is the key cytotoxic event in these cells, we assessed the effect of combined p97 inhibitor CB-5083/PARPi talazoparib exposure in DLD1 cells with/without engineered genetic ablation of the *BRCA2* gene (DLD1 and DLD1.*BRCA2^−/−^*). CB-5083, when used alone, had a modest *BRCA2* synthetic lethal effect (Supplementary Figure 6C) but adding CB-5083 to the PARPi talazoparib had a far greater effect on cell inhibition in DLD1 *BRCA2^−/−^* cells than in isogenic cells with wild-type *BRCA2* (Figure 6F and Supplementary Figure 6D, E). To assess these synthetic lethal effects in the setting of a *BRCA1* defect, we used tumour-derived organoids. In the first instance, we used tumour organoids derived from mice with combined *Brca1/Tp53* loss-of-function mutations (WB1P^36^) and found that CB-5083 further sensitised tumour organoids to the talazoparib (Figure 6G and Supplementary Figure 6F). For example, 100 nM CB-5083 enhanced the organoid inhibitory effect of 6 pM talazoparib, a PARPi concentration that when used as a single-agent, had no detectable effect on cell survival. Secondly, we assessed p97 inhibitor CB-5083/PARP inhibitor talazoparib combinations in a human patient-derived tumour organoid model (PDO) derived from a triple negative breast cancer patient who harboured a deleterious *BRCA1* mutation (KCL014BCPO), derived from a tumour from a patient with a germline pathogenic BRCA1 p.R1203* mutation (*BRCA1* c.3726C>T) which was homozygous in the organoid. Similar to the mouse tumour organoid, CB-5083 led to marked shift in talazoparib sensitivity (Figure 6H and Supplementary Figure 6G), suggesting that p97 inhibition has the potential to potentiate the effects of PARPi in human tumour cells.

## Discussion

The effectiveness of PARPi when used in the treatment of cancer relies at least in part upon both the ability of these drugs to trap PARP1 in the chromatin fraction combined with the inability of some tumour cells to repair the resultant DNA lesions by processes such as homologous recombination. Understanding how trapped PARP1 is removed from the chromatin fraction could have the potential to lead to greater insight as to how PARPi could be best used. Here, by identifying proteins that associate with trapped PARP1 we elucidated a biochemical cascade that processes trapped PARP1. In this pathway trapped PARP1 is sequentially SUMOylated by PIAS4 and ubiquitylated by RNF4. The RNF4-dependent poly-K48-ubiquitin chains on trapped PARP1 facilitates p97 binding via UFD1. Once recruited, p97 ATPase activity is necessary to drive PARP1 eviction from the chromatin (Figure 6I). Of note, recently two other factors – the E3 ubiquitin ligase TRIP12^37^ and the deubiquitinylating enzyme ATXN3^38^ were demonstrated to effect the dynamics of PARP1. Importantly, both factors are recruited to PARP1 in a PAR-dependent manner. Under trapping conditions used in this work, PAR is absent (for example Figure 4B, H), and it is the SUMO/ubiquitin (Figure 2-4) that drives the signals for the p97-dependent axis to operate and modulate the inhibitor-induced trapped complex. Interference with any of the trapped PARP1 processing steps ultimately leads to persistence of the trapped complex. Consistent with both PARP1 trapping being a key determinant of PARPi-induced cytotoxicity and p97 playing a key role in removing trapped PARP1, p97 inhibitors enhance the tumour cell inhibitory effects of PARPi, not only in isogenic cell lines with *BRCA*-gene defects, but also in tumour organoids, including those derived from a breast cancer patient with a *BRCA1* mutation. By extension, the ability to modulate the release of PARP1 from DNA after trapping offers the possibility to optimise and augment the application of PARPi.

We also see a number of new questions that might now arise from these observations. Firstly, although PIAS4 and RNF4 appear to act in a linear manner, there remains the possibility that the balance of SUMOylation and ubiquitylation is influenced by other E3 ligases. Indeed, we observed residual modifications in the knock out models that we have used and the effect on trapped PARP1 resolution is modest for the *PIAS4^-/-^* cells (Figure 5C). Intriguingly, we noted that the decreased ubiquitylation in *RNF4^−/−^* cells is concomitant with increased SUMOylation. Regarding p97 recruitment, our data suggest that UFD1 is required for the recruitment of p97 to trapped PARP1 (Figure 4J). How exactly UFD1 recruits p97 to trapped PARP1 remains to be established. UFD1 is a well-known ubiquitin-chain reader as it possesses Ub-binding domain. In yeast, UFD1 has been shown to bind SUMO (in addition to ubiquitin) and to recruit p97/cdc48 to SUMOylated substrates^39, 40^. However, UFD1 binding to SUMO has never been demonstrated in mammalian cells. Our data presented here, suggests that p97 recruitment to trapped PARP1 depends on RNF4-dependent ubiquitylation; it thus seems likely that UFD1 recruits p97 via its canonical role as an ubiquitin-chain reader, directly bridging p97 and the ubiquitin chains on p97 substrates, in this case ubiquitylated PARP1. Secondly, there are other known modulators of PARP1 trapping. Recent observations suggest that altering the levels of this residual PAR, for example by altering the activity of the PAR-glycohydrase, PARG, alters PARPi-induced PARP1 trapping^5^. How, or indeed whether, p97 activity on trapped PARP1 is modulated by residual PAR and PARG remains to be determined, as does whether PARG and p97 work independently upon trapped PARP1 or in a more co-ordinated fashion.

Finally, we also highlight that the PARP inhibitors-generated DNA lesions, appear to be processed via methods somewhat analogous to those caused by Topoisomerase I inhibitors, namely trapped TOP1-cleavage complexes^10^. For example, both PARPi and TOP1 inhibitors cause replication fork stress and sensitivity in cells with homologous recombination defects and the sensitivity to both classes of agents is also modulated by *SLFN11* status^41^. Although trapped TOP1-cleavage complexes form a covalent link with DNA, and as far as we are aware, trapped PARP1 does not, both are SUMOylated, ubiquitylated and modified by p97 (reviewed in^42^ and data shown here). Given these similarities, one might imagine that the processes that activate the SUMOylation and ubiquitylation of trapped PARP1 and trapped TOP1 and TOP2 cleavage complexes might also be shared and not necessarily private to the precise nature of the nucleoprotein complexes that they cause, but possibly more to do with their ability to interfere with the normal progression of DNA metabolic processes including replication and transcription. One might therefore speculate that the SUMOylation of trapped PARP1 might be controlled and instigated by the stalling of replication forks rather than the direct detection of trapped PARP1 in chromatin *per se*.

In conclusion, the work described here elucidates an elegant and highly orchestrated molecular machinery, composed of the E3-SUMO ligase PIAS4, the SUMO-chain reader and ubiquitin ligase RNF4, the ubiquitin chain reader UFD1 and the ATPase p97. Together, these recognise and remove trapped PARP1 from chromatin.

## Materials and Methods

### Cells and cell culture

CAL51 and DLD1 cells were maintained in Dulbecco’s modified Eagle’s medium (DMEM) supplemented with 10 % FBS, 1x Penicillin-Streptomicyn (Sigma-Aldrich). CAL51 PARP1-/-cells were previously described^14^. They were transfected with a corresponding PARP1-expressing piggyBac construct in combination with High-PBase-expressing plasmid^43^. 72 hours after transfection, single-cell clones were sorted by FACS and allowed to expand. These clones were characterized for the expression of the tagged protein by microscopy and western blotting. HEK293 *PARP1-/-* cells were a kind gift from Ivan Ahel, University of Oxford. The HCT116 *PIAS4^−/−^* and MCF7 *RNF4^−/−^* cells were previously described^22^. WB1P organoid line was previously described ^44^. They were grown in a mix of 50 % Matrigel (Corning) and 50 % Advanced DMEM/F12 (Life Technologies) containing 10 mM HEPES (Sigma-Aldrich) pH 7.5, GlutaMAX (Invitrogen) and supplemented with 125 mM N-acetylcysteine (Sigma-Aldrich), B27 supplement and 50 ng/ml EGF (Life Technologies). 24 hours prior to drug organoids were seeded at 10000 cells per well of a 24 well plate and drugs added at the indicated concentrations. Cell viability was assessed using 3D cell-titer glow (Promega). KCL014BCPO was derived (Badder et al., manuscript in preparation), similarly to as described^45^. Briefly, human breast tumour samples were obtained from adult female patients after informed consent as part of a non-interventional clinical trial (BTBC study REC no.: 13/LO/1248, IRAS ID 131133; principal investigator: A.N.J.T., study title: ‘Analysis of functional immune cell stroma and malignant cell interactions in breast cancer in order to discover and develop diagnostics and therapies in breast cancer subtypes’). This study had local research ethics committee approval and was conducted adhering to the principles of the Declaration of Helsinki. Specimens were collected from surgery and transported immediately. A clinician histopathologist or pathology-trained technician identified and collected tumour material into basal culture medium. Tumour samples were coarsely minced with scalpels and then dissociated using a Gentle MACS dissociator (Miltenyi). The resulting cell suspension was mechanically disrupted, filtered and centrifuged. Resulting cell pellets were then plated into 3D cultures at approximately 1 × 10^3^ to 2 × 10^3^ cells per μl in Ocello PDX medium (OcellO B.V) and hydrogel. All cultures were maintained in humidified incubators at 37 °C, 5 % CO_2_. All human cell line identities were confirmed by STR typing and verified free of mycoplasma infection using Lonza MycoAlert.

### Plasmids, antibodies and reagents

PB-PARP1-eGFP – PARP1 cDNA was cloned in a previously described piggyBac vector^46^. To generate PARP1-Apex2-eGFP construct, the *Apex2* gene was amplified from Addgene vector 49386 and inserted inbetween PARP1 and eGFP coding sequences via InFusion (Clonetek, 648910). PBZ-mRuby2 is described in^14^. UB-STREP-HA was a kind gift from Vincenzo D’angiolella, HA-SUMO2^47^, FLAG-PARP1 was a kind gift from Ivan Ahel, p97-GFP was a kind gift from Hemmo Mayer.

The wild-type PIAS4 expressing construct was obtained from Addgene (#15208) and RNF4 from Origene (RC207273). The corresponding SAP and SIM mutants were generated as described^22^. The RNF4-M136S,R177A was a kind gift from Ronald Hay.

Antibodies used were: GFP (Sigma-Aldrich, 11814460001); PARP (CST, 9532) for immunoblotting and PLA; p97 (Abcam, ab11433) for immunoblotting and PLA; PAR (Trevigen, 4335-AMC-050); HA (Roche, 11867423001); FLAG (M2, Sigma-Aldrich, F1804) for immunoprecipitation; FLAG (Sigma-Aldrich F7425) for immunoblotting; Streptavidin-HRP (ThermoFisher, S911); PARP1 (Sigma-Aldrich, WH0000142M1) for PLA; β-actin (Invitrogen, AM4302); lamin-B1 (Thermo, PA5-19468); vinculin (Abcam, ab18058); phospho-H2AX (CST, 9718S) for PLA; phospho-H2AX (Millipore, 05-636) for foci immunostaining; RAD51 (Abcam, ab133534) for foci immunostaining; Histone H3 (CST, 9715); SUMO1 (CST, 4940); SUMO2/3 (CST, 4971); ubiquitin (Santa Cruz Biotechnology, sc-8017); RNF4 (Novusbio, NBP2-13243); UFD1L (Abcam, ab181080); Anti-Rabbit IgG HRP (Rockland, 18-8816-31). Talazoparib was supplied by Pfizer as part of the BCN Catalyst programme. Other small molecules were as follows: Olaparib (Selleckchem, S1060); Veliparib (Selleckchem, S1004); UKTT15 from in-house synthesis as described in^4^, MMS (Sigma-Aldrich, 129925-5G); CB-5083 (Selleckchem, S8101); CuET from in-house synthesis as described in^33^; MLN-7243 (Selleckchem, S8341); ML-792 (Medchemexpress, HY-108702). siRNAs were obtained from Dharmacon: RNF4 (L-006557-00-0005 and 3’UTR siRNA sequence 5’-GGGCAUGAAAGGUUGAGAAUU); UFD1L (L-017918-00-0005); NPL4 (L-020796-01-0005).

### Western blotting

Standard protocols for sodium dodecyl sulfate-polyacrylamide gel electrophoresis (SDS PAGE) and immunoblotting were used ^48^. Nitrocellulose membrane (GEHealthcare) or PVDF (BioRad) were used to transfer proteins from polyacrylamide gels depending on the antibody.

### Cellular fractionation immunoprecipitation

Cells were washed two times with PBS then resuspended in buffer A (10 mM HEPES, 10 mM KCI, 340 mM sucrose, 10 % glycerol, 2 mM EDTA, protease and phosphatase inhibitors, N-ethylmaleimide (NEM)). Triton X-100 was added to achieve a final concentration of 0.1 % and left on ice for 2-5 min depending on cell line. The supernatant was harvested as the cytosolic fraction and the pellet (nuclei) was then washed two times with buffer A. Buffer B (3 mM EDTA, 0.2 mM EGTA, 5 mM HEPES pH 7.9, protease and phosphatase inhibitors, NEM) was then added to burst nuclei, after which lysates were kept on ice for 10 min. Supernatant was then removed as the nuclear soluble fraction. Remaining chromatin pellet was then washed with buffer B in 0.5 % Triton X-100 followed by benzonase buffer without MgCl_2_ (50 mM Tris-HCl pH 7.9, 100 mM NaCl). Benzonase digestion buffer (50 mM Tris-HCl pH7.9, 10 mM MgCl_2_, 100 mM NaCl, protease and phosphatase inhibitors, NEM) supplemented with 125 U of Benzonase enzyme (Merk Millipore) and rotated on wheel at 4°C for 1 hour. Samples were then centrifuged at 20,000 g for 15 min, chromatin input for the immunoprecipitation reaction was then taken from supernatant. The remaining supernatant was then incubated with respective beads on a rotating wheel for 3 hours at 4°C. For native IPs, 1:200 ethidium bromide was added to remove unwanted DNA-protein interactions. Beads were then washed with IP wash buffer (50 mM Tris-HCl pH 7.4, 150 mM NaCl, 0.5 mM EDTA, 0.05 % Triton X-100) three times before elution with lamelli buffer.

### Whole cell Immunoprecipitation

Cells were lysed in IP lysis buffer (50 mM Tris HCl pH 7.4, 150 mM NaCl, 1 mM EDTA, 0.5 % Triton X-100, protease and phosphatase inhibitors, NEM) and spun on wheel at 4°C for 10 min. The supernatant was removed and pellet was washed with benzonase buffer once and the supernatants were pooled together. Benzonase buffer (50 mM Tris-HCl pH 7.9, 10 mM MgCl_2_, 100 mM NaCl, protease and phosphatase inhibitors, NEM) supplemented with 125 U of Benzonase enzyme (Merk Millipore) was added to the pellet and left on wheel at 4°C for 1 hour. Samples were then centrifuged at 20,000 g and all supernatants were pooled. Input for the IP was then removed and samples were incubated with respective beads on wheel for 3 hours at 4°C. Beads were then washed with IP wash buffer (50 mM Tris-HCl pH 7.4, 150 mM NaCl, 0.5 mM EDTA, 0.05 % Triton X-100) three times before elution with laemlli buffer.

### Denaturing IP

Cells were lysed either according to fractionation immunoprecipitation or whole cell immunoprecipitation protocol as stated above. Before incubation with beads, SDS was added to samples to a concentration of 1 % and boiled at 95°C for 5 min. Samples were then diluted in 1 % Triton X-100 to achieve a dilution of 1:10 (SDS at 0.1 %) along with beads and rotated on wheel at 4°C for 3 hours. Beads were then washed with IP wash buffer (50 mM Tris-HCl pH 7.4, 150 mM NaCl, 0.5 mM EDTA, 0.05 % Triton X-100) three times before elution with laemlli buffer.

### Cell viability and clonogenic survival assays

The viability of cells was measured after six days exposure to various concentrations of drugs using the Cell Titre-Glo assay (Promega). Long-term drug exposure effects were assessed by colony formation assay after 12–14 days exposure to a drug-containing medium (refreshed weekly) and cells stained at the end of the assay with sulforhodamine B. When plotting survival curves, the surviving fraction was calculated relative to DMSO (solvent)-exposed cells.

The viability of the KCL014BCPO organoid line was measured using the 3D Cell Titre-Glo assay (Promega). Organoids were seeded in 24-well plates, with one 15 µl Matrigel droplet containing 3000 cells per well. 24 hours after seeding, the organoids were treated with a drug-containing media (drug refreshed after 4 days) for 7 days, before assessing the viability by measuring 3D Cell Titre-Glo luminescence.

### Apex2-mediated Proximity Labelling

For each condition tested, 5-10 x 10^6^ cells expressing PARP1-Apex2-eGFP were exposed to either 0.01 % MSS or a combination of 0.01 % MSS + 100 nM talazoparib for 1 hour. In the last 30 min of the incubation, 500 µM final concentration biotin-tyramide (Sigma-Aldrich, SML2135) was added to the media. To label proteins, 1 mM final concentration H_2_O_2_ (Sigma-Aldrich, H1009) was added for 60 s. The reaction was quenched by washing the cells tree times with freshly prepared quench solution (PBS containing 10 mM Sodium ascorbate, 10 mM Sodium Azide, 5 mM Trolox (Sigma-Aldrich, 238813)). Subsequently, the cells were scraped in quench solution and washed twice in 0.1% IGEPAL CA-630 quench solution to remove the fraction of cytosolic proteins. The remaining nuclei were lysed in nuclear RIPA buffer (50 mM TrisHCl pH 7.5, 1 M NaCl, 1 % IGEPAL CA-630, 0.1 % sodium deoxycholate, 1 mM EDTA) for 10 min on ice. The lysates were diluted with RIPA buffer not containing NaCl to final 200 mM NaCl, sonicated for 1 min and incubated with 250 U benzonase for 20 min at room temperature.

The lysates were clarified by centrifugation at 13000 g for 15 min at 4DC. Protein concentration was evaluated and 1 mg total protein was incubated with 30 μl streptavidin-magnetic beads (ThermoFisher, 88816) for 1 h at room temperature. The beads were washed stringently by sequential washes: twice with RIPA lysis buffer, once with 1 M KCl, once with 0.1 M Na_2_CO_3_, once with 2 M urea, twice with RIPA lysis buffer and processed further for MS analysis.

### Rapid immunoprecipitation mass spectrometry of tagged protein (RIME-based approach)

For each condition tested, 5-10 x 10^6^ cells were exposed to either 0.01% MSS or a combination of 0.01% MSS + 100 nM talazoparib for 1 hour. At the end of the incubation period, formaldehyde (ThermoFisher, 28908) was added to the media to 1 % final concentration and incubated for 10 min at room temperature. The reaction was quenched by the addition of 125 mM glycine final concentration. The cells were collected and washed once in ice-cold PBS. The cells were resuspended in ice cold PBS containing 0.1 % Triton X-100 and protease inhibitor cocktail (Merck, 4693116001), in order to extract the cytoplasm. The nuclei were spun at 3000 g for 5 min at 4°C, and subsequently resuspended in PBS containing 1 % IGEPAL CA-630 (Sigma-Aldrich) and protease inhibitors, and incubated on ice for 15 min in order to release the nuclear soluble proteins. The remaining chromatin was spun at 13000 g for 5 min at 4°C and resuspended in PBS containing 0.1 % IGEPAL CA-630 and protease inhibitors. The chromatin pellet was spun at 13000 g for 5 min at 4°C, resuspended in lysis buffer (20 mM HEPES pH 7.5, 150 mM NaCl, 0.5 % sodium deoxycholate, 0.1 % SDS, 10 mM MgCl_2_) supplemented with 250 U benzonase (Sigma-Aldrich, E1014) and incubated for 30 min at room temperature with rotation to release the chromatin bound proteins. The supernatant was isolated after centrifugation (13000x g for 10 min at 4°C) and incubated with 25 μl GFP Trap (Chromotek, gtm-20) magnetic beads for 1 hour at 4°C with rotation. The beads were washed four times with the lysis buffer and processed further for MS analysis.

### Mass spectrometry and data analysis

After initial washes according to the purification method, the beads were further washed twice with 50 mM ammonium bicarbonate. The proteins on the beads were digested with 0.1 μg/μl sequencing grade trypsin (Roche) overnight at 37°C. The peptide solution was neutralised with 5% formic acid, acetonitrile was added to 60% final concentration and the solution was filtered through a Millipore Mutiscreen HTS plate (pre washed with 60 % acetonitrile). The peptide solution was lyophilised on a SpeedVac and the peptides were dissolved in 20 mM TCEP-0.5 % formic acid solution. The LC-MS/MS analysis was conducted on the Orbitrap Fusion Tribrid mass spectrometer coupled with U3000 RSLCnano UHPLC system (ThermoFisher). The peptides were first loaded on a PepMap C18 trap (100 μm i.d. x 20 mm, 100 Å, 5 μm) at 10 μl/min with 0.1% FA/H_2_O, and then separated on a PepMap C18 column (75 µm i.d. x 500 mm, 100 Å, 2 μm) at 300 nl/min and a linear gradient of 4-32 % ACN/0.1 % FA in 90 min with the cycle at 120 min. Briefly, the Orbitrap full MS survey scan was m/z 375–1500 with the resolution 120,000 at m/z 200, with AGC (Automatic Gain Control) set at 40,000 and maximum injection time at 50 ms. Multiply charged ions (z = 2 – 5) with intensity above 8,000 counts (for Lumos) or 10,000 counts (for Fusion) were fragmented in HCD (higher collision dissociation) cell at 30 % collision energy, and the isolation window at 1.6 Th. The fragment ions were detected in ion trap with AGC at 10,000 and 35 ms maximum injection time. The dynamic exclusion time was set at 40 s with ±10 ppm.

The mass spectrometry proteomics data have been deposited to the ProteomeXchange Consortium via the PRIDE^49^ partner repository with the dataset identifier PXD024337. Raw mass spectrometry data files were analysed with Proteome Discoverer 1.4 (Thermo). Database searches were carried out using Mascot (version 2.4) against the Uniprot human reference database (January 2018, 21123 sequences) with the following parameters: Trypsin was set as digestion mode with a maximum of two missed cleavages allowed. Precursor mass tolerance was set to 10 ppm, and fragment mass tolerance set to 0.5 Da. Acetylation at the N terminus, oxidation of methionine, carbamidomethylation of cysteine, and deamidation of asparagine and glutamine were set as variable modifications. Peptide identifications were set at 1 % FDR using Mascot Percolator. Protein identification required at least one peptide with a minimum score of 20. For the Apex2-based proximity labelling MS the following steps were taken. Proteins identified with a single peptide were removed from further analysis. PARP1-eGFP MS profile under trapping conditions (MMS + talazoparib) was used as a negative control. Proteins identified with >2 unique peptides in this sample were removed from further analysis. Peptide Spectrum Matches (PSM) was used as a proxy of protein abundance in the samples. A ratio was built between PARP1-Apex2-eGFP MMS+talazoparib PSM and PARP1-Apex2-eGFP MMS PSM as an indicator for enrichment in the trapping conditions. Where PSM values were absent from the PARP1-Apex2-eGFP MMS PSM (i.e. no detection in the sample) a value of one was added in order to calculate a meaningful ratio (the data is provided in Supplementary Table 1). The list of genes was then searched on the STRING database to build a network of the hits. A high confidence threshold was set for mapping the network, using a minimum required interaction score of 0.7 for connecting nodes. Single, unconnected nodes were excluded from the network plots. The gene list was searched in the Enrichr database to assess which KEGG 2019 pathway annotations are enriched in the dataset. The list of annotations was filtered using -log(p) values of 1.3 (p = 0.05) or 2 (p = 0.01) (the data is provided in Supplementary Table 2). For RIME analysis the following steps were taken. Proteins identified with single peptides were removed from further analysis. Proteins identified in CAL51 *PARP1^−/−^* cells were considered as background and removed from further analysis when they were identified with more than two unique peptides. Subsequently the MS data obtained from PARP1^WT^-eGFP or PARP1^del.p.119K120S^-eGFP cells were considered separately. For each cell line a ratio was built between the MMS+talazoparib PSM and the MMS PSM as an indicator for enrichment in the trapping conditions. Where PSM values were absent from the MMS PSM (i.e. no detection in the sample) a value of one was added in order to calculate a meaningful ratio (the data is provided in Supplementary Table 3 and 4 for PARP1^WT^-eGFP or PARP1^del.p.119K120S^-eGFP, respectively).

### Proximity ligation assay (PLA)

Proximity ligation assays were carried out with Duolink^®^ In Situ Red Starter Kit Mouse/Rabbit (Sigma-Aldrich) according to the manufacturer’s protocol. The primary antibodies used were mouse anti-PARP1 (WH0000142M1-100UG), rabbit anti-PARP (Cell Signalling), mouse anti-p97 (ab11433) and rabbit anti-phospho-H2AX (Cell Signalling). The antibodies were used in 1:1500 dilution. Images were acquired on Marianas advanced spinning disk confocal microscope (3i) and analysed with a custom CellProfiler pipeline. Typically, several hundred nuclei were counted per condition from at least two independent biological repeats.

### Microirradiation

Cells were grown in glass-bottom culture dishes (MaTek,P35G-0.170-14-C) in 10% FBS DMEM media and maintained at 37 °C and 5%CO2 in an incubation chamber mounted on the microscope. Imaging was carried out on Andor Revolution system, Å∼60 water objective with micropoint at 365 nm. For FRAP analysis the cells were acquired one at a time; each cell was irradiated at a single spot with 1 µm diameter within the nucleus. After the signal of recruitment reached its maximum (typically 30 s to 60 s after microirradiation) the recruitment spot was bleached with a 488 nm laser and imaging continued with one frame per 2 s intervals. For each experimental condition ten to twelve cells were acquired and the experiment was repeated independently on a different imaging day. From the raw intensities of the microirradiation site, the spot intensity immediately prior to bleaching was set to 1 and immediately after bleaching to 0. The recovery data was fitted with one site-specific binding model of non-linear regression (Graphpad Prism software) and the extra sum of squares *F* test was used to calculate the *t_1/2_*.

### Cell cycle analysis

Cells were incubated in the presence of inhibitors for the corresponding amount of time. One hour prior to fixation 10 µM ethylene-deoxyuridine (EdU, Thermofisher) was added to the media. Subsequently, the cells were trypsinized and fixed in ice-cold absolute ethanol. The cells were re-hydrated via PBS wash and permeabilized with 0.5 % Triton X-100 in PBS for 15 min at room temperature with rotation. After a PBS wash, a click chemistry reaction cocktail was added to the cells (100 mM Tris HCl pH 7.6, 4 mM CuSO_4_, 2.5 µM azide-fluor488 (Sigma), 100 mM Sodium ascorbate (Sigma)) and incubated for 30 min at room temperature, protected from light. After a PBS wash, propidium iodide/RNase staining solution (Thermo) was added to the cells for 30 min. The cell cycle profiles were acquired on a BD LSRII flow cytometer and analysed with the BD FACSdiva software.

### Chromatin fractionation

The chromatin fractionation assay for PARP trapping was based on a previously published protocol^2^. For the trap-chase experiments cells were grown in six-well plates, exposed to 100 nM talazoparib and 0.01% MMS for 1 h and subsequently incubated in a media containing the corresponding drugs (typically 100 nM talazoparib, 10 μM CB-5083, 1 μM CuET) for a chase period of 3 h. The cells were fractionated with the Subcellular Protein Fractionation kit for Cultured Cells (ThermoFisher #78840) according to the manufacturer’s recommendations.

## Acknowledgements

We thank all the members of the Lord, Tutt and Ramadan laboratories for the useful discussion of this project. This work was funded by Cancer Research UK, as part of Programme Grant funding to CJL, by Breast Cancer Now as part of Programme Funding to the Breast Cancer Now Toby Robins Research Centre (CJL, ANJT, SP), by Breast Cancer Now Catalyst Funding (CJL) and by Medical Research Council Programme Grant (MC_PC 12001/1 and MC_UU 00001/1, KR) and Breast Cancer Now Project funding (2019DecPR1406, KR). JB was support by the Danish Cancer Society (R204-A12617-B153), and the Swedish Cancerfonden ( #17017). Organoid line derivation in AT laboratory was funded by NC3Rs funding to AT and CJL (NC/P001262/1) and we also thank Breast Cancer Now, working in partnership with Walk the Walk, for supporting the work of Patient Derived Models Team at the Breast Cancer Now Toby Robins Research Centre. This work represents independent research supported by the National Institute for Health Research (NIHR) Biomedical Research Centre at The Royal Marsden NHS Foundation Trust and the Institute of Cancer Research, London. The views expressed are those of the authors and not necessarily those of the NIHR or the Department of Health and Social Care. Y.P. and Y.S. are supported by the Center for Cancer Research, the Intramural Program of the National Cancer Institute, NIH (Z01-BC 006150).

## Supplementary Figures

**Supplementary Figure 1.**
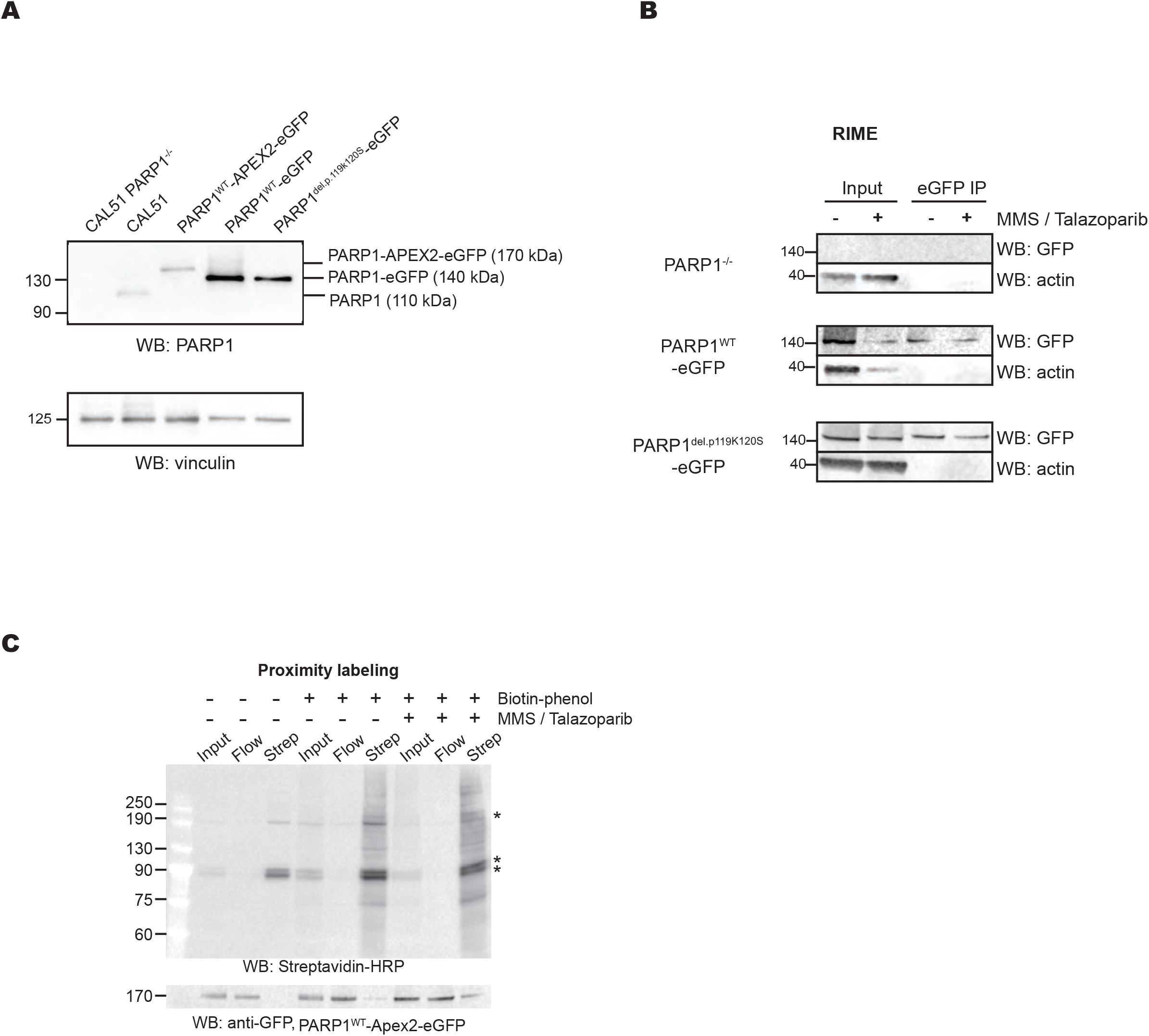
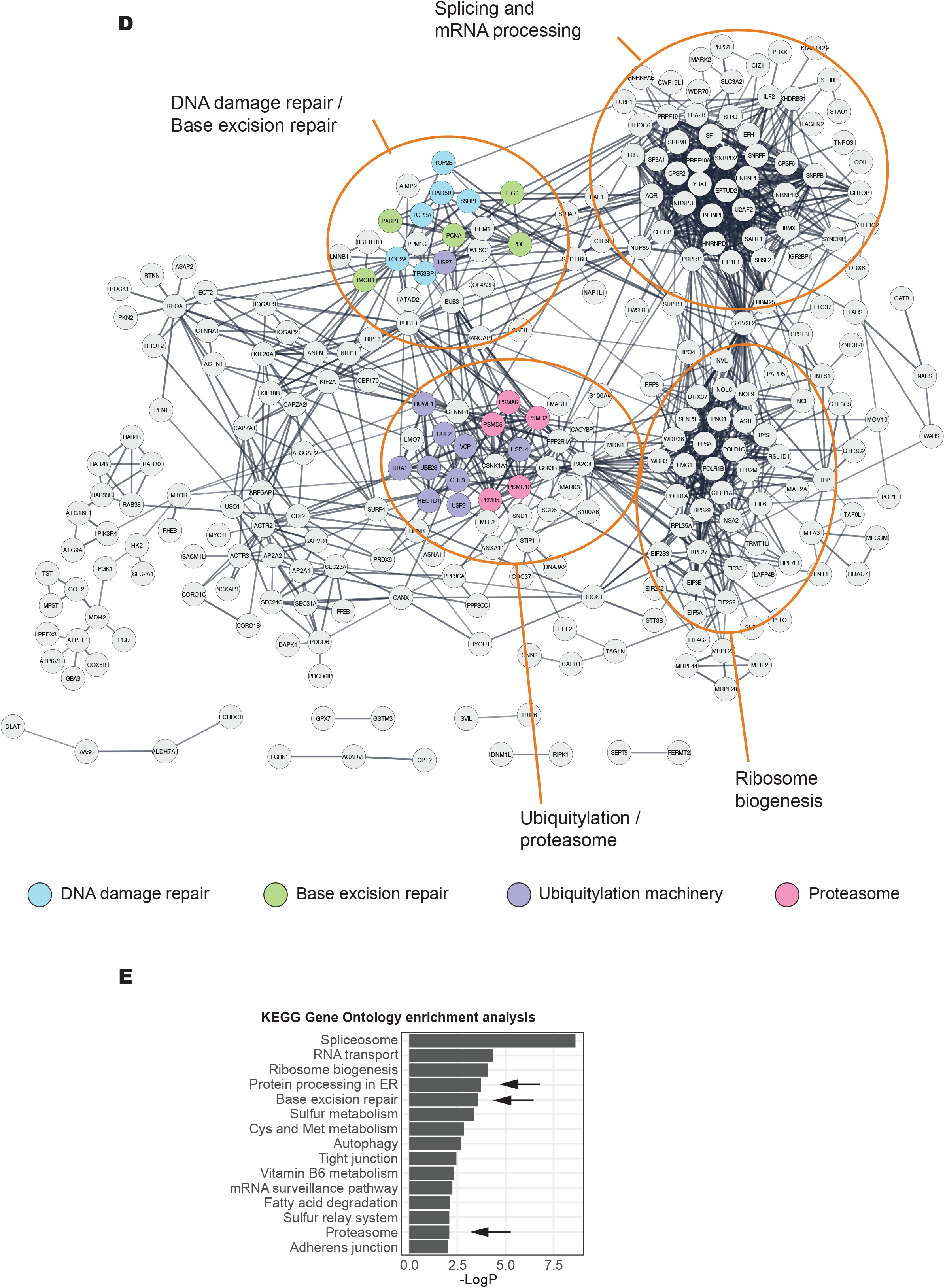
**A.** A Western blotting for the expression of the tagged PARP1, detected by an PARP1 antibody. Three single-cell clones, chosen for this study (PARP1*^WT^*-eGFP, PARP1*^del.p.119K120S^*-eGFP and PARP1-Apex2-eGFP) show the expression of the tagged protein at the expected molecular weight. **B.** A Western blot analysis of the purified PARP1-associated proteins as described in the RIME experiment in Figure 1A. **C**. Western blot analysis of the purified biotinylated proteins isolated in the PARP1WT-Apex2-eGFP proximity labelling experiment. Immunoblotting using Streptavadin-HRP is shown in the top panel, whilst anti-GFP immunoblotting is shown in the bottom panel. Endogenously biotinylated proteins are indicated as*. **D**. STRING network diagram of proteins identified by PARP1 proximity labelling under PARPi trapping conditions (as described in Methods). The graph shows connected nodes identified with a high stringency threshold of 0.7 (non-connected proteins are excluded from this visualisation). The colour coding corresponds to the following functional annotation: DNA damage repair-associated proteins (blue), base excision repair (green), ubiquitylation machinery (purple) and proteasome (magenta). Clusters, enriched for certain biological processes are indicated (e.g. “Ubiquitylation/proteosome”). **E**. Summary of gene set ontology analysis of the networks presented in (F). KEGG terms enriched at p-value <0.01 are shown.

**Supplementary Figure 2.**
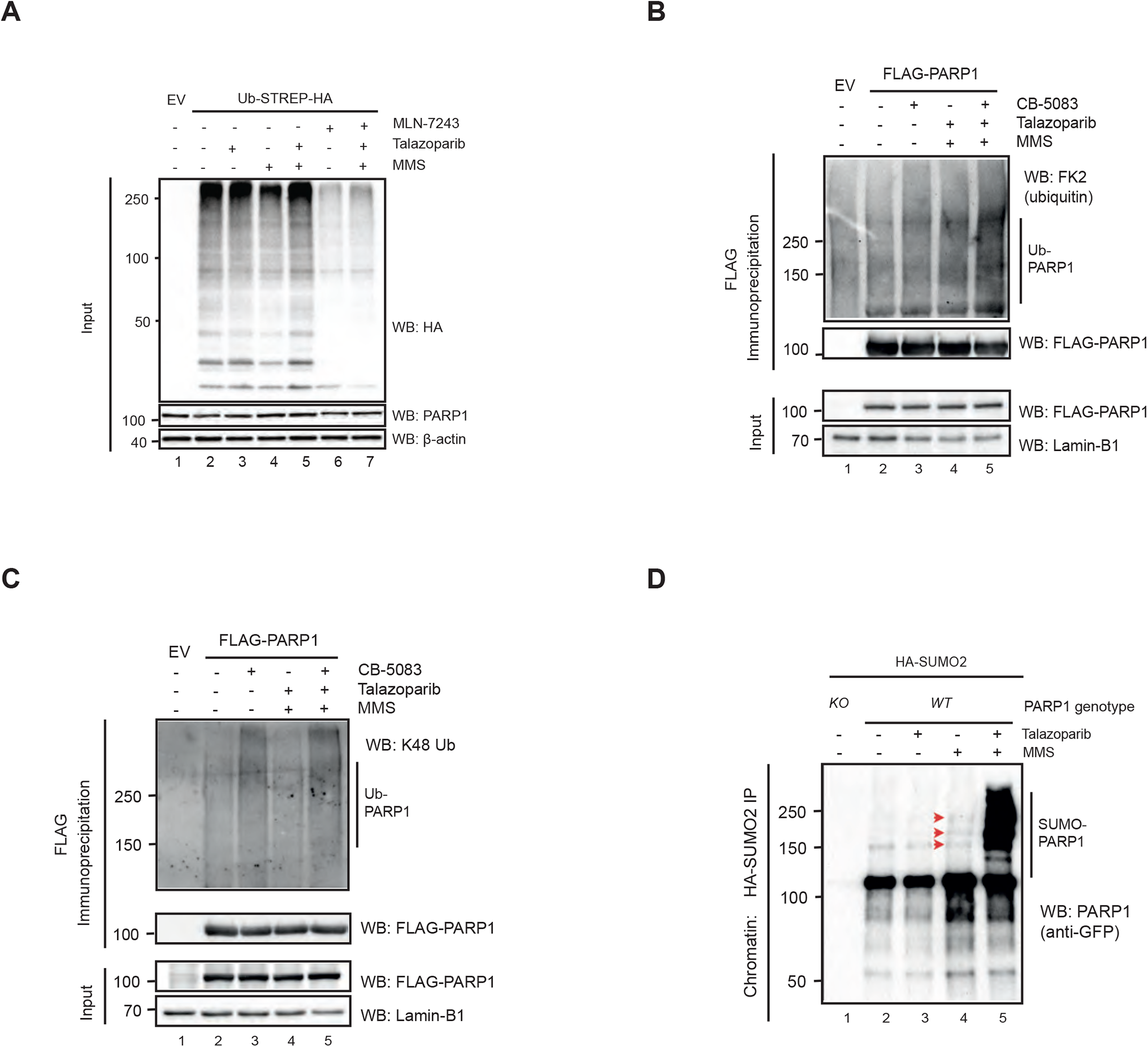
**A.** Input controls for Figure 2C, showing the efficacy of MLN-7243 to inhibit ubiquitylation. **B.** High exposure blot of PARP1 SUMOylation from Figure 2D, red arrows show SUMOylated PARP1 in MMS treated samples. **C.** Reciprocal denaturing IP over PARP1-FLAG showed accumulation of trapped PARP1 ubiquitination in HEK293s. HEK293 cells were transfected with PARP1-FLAG-expressing construct for 24 hours then treated with 100 nM talazoparib/0.01 % MMS and/or 10 µM CB-5083. Cells were lysed, chromatin was digested and then incubated with anti-FLAG beads. 4 % of sample was harvested for input pre-incubation. **D.** As in (C), but the immunoprecipitated proteins were analysed with an anti-K48 Ub chains recognising antibody.

**Supplementary Figure 3.**
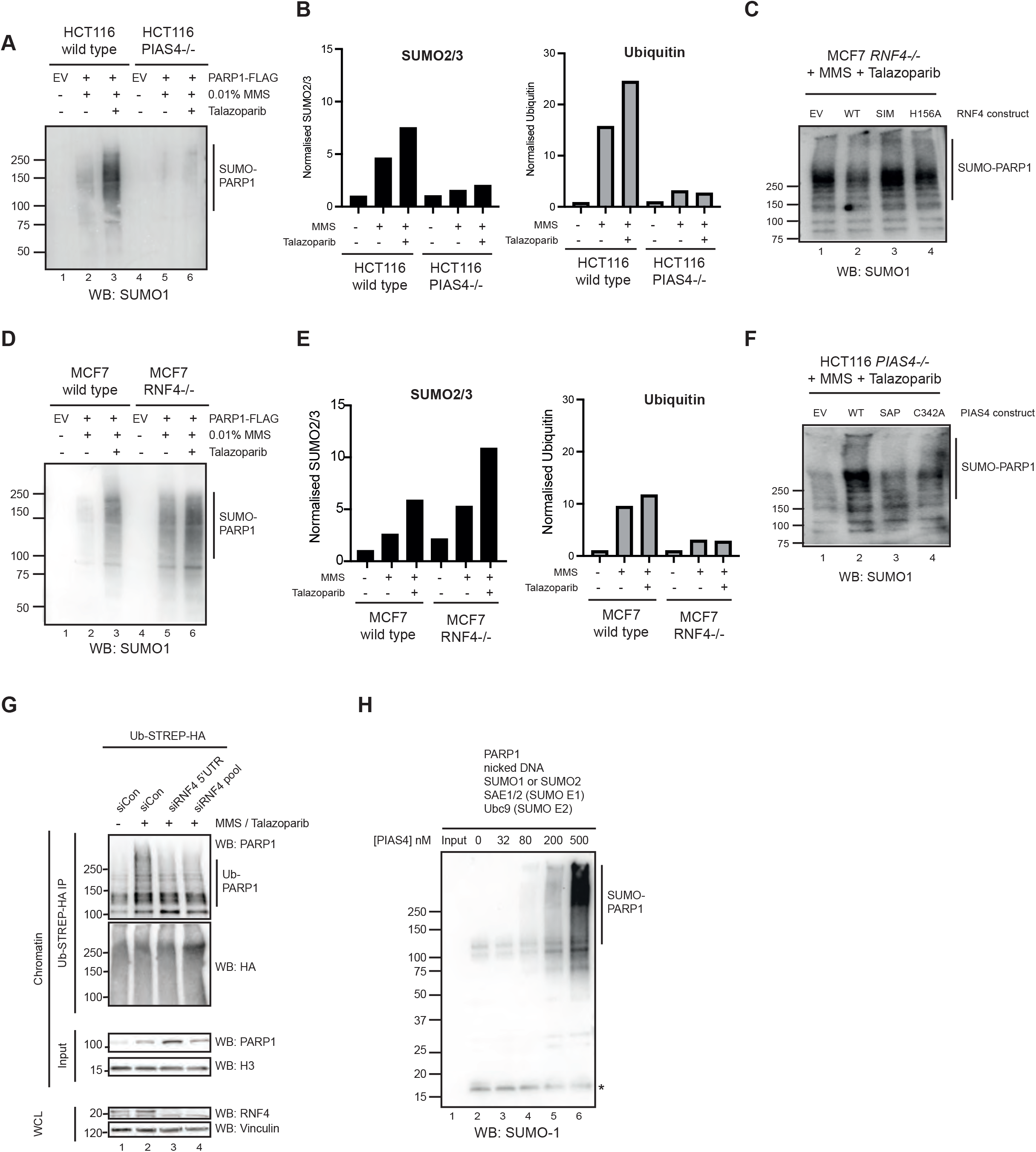
**A**. PARP1 SUMOylation by SUMO1 was detected as described in Figure 3A. **B**. A quantification of the SUMO2/3ylated and ubiquitylated PARP1 isoforms in the gels in Figure 3A. **C**. PARP1 SUMOylation by SUMO1 detected as described in Figure 3B. **D**. PARP1 SUMOylation by SUMO1 was detected as described in Figure 3D. **E**. A quantification of the SUMO2/3ylated and ubiquitylated PARP1 isoforms in the gels in Figure 3D. **F**. PARP1 SUMOylation by SUMO1 detected as described in Figure 3E. **G.** RNF4 depletion prevents PARP1 ubiquitination under PARP trapping conditions. Denaturing IP of UB-HA-STREP-expressing HEK293 cells, similar to Figure 2B. Cells were depleted of RNF4 with either a 5’UTR sequence or Dharmarcon SMARTpool and were treated with 100 nM Talazoparib and 0.01 % MMS. Pulldown was conducted with Streptactin beads. **H**. *In vitro* SUMOylation assay as described in Figure 3G. The reactions were incubated in the presence of SUMO1, which was subsequently detected by anti-SUMO1 antibody. The asterisk indicates the free SUMO1.

**Supplementary Figure 4.**
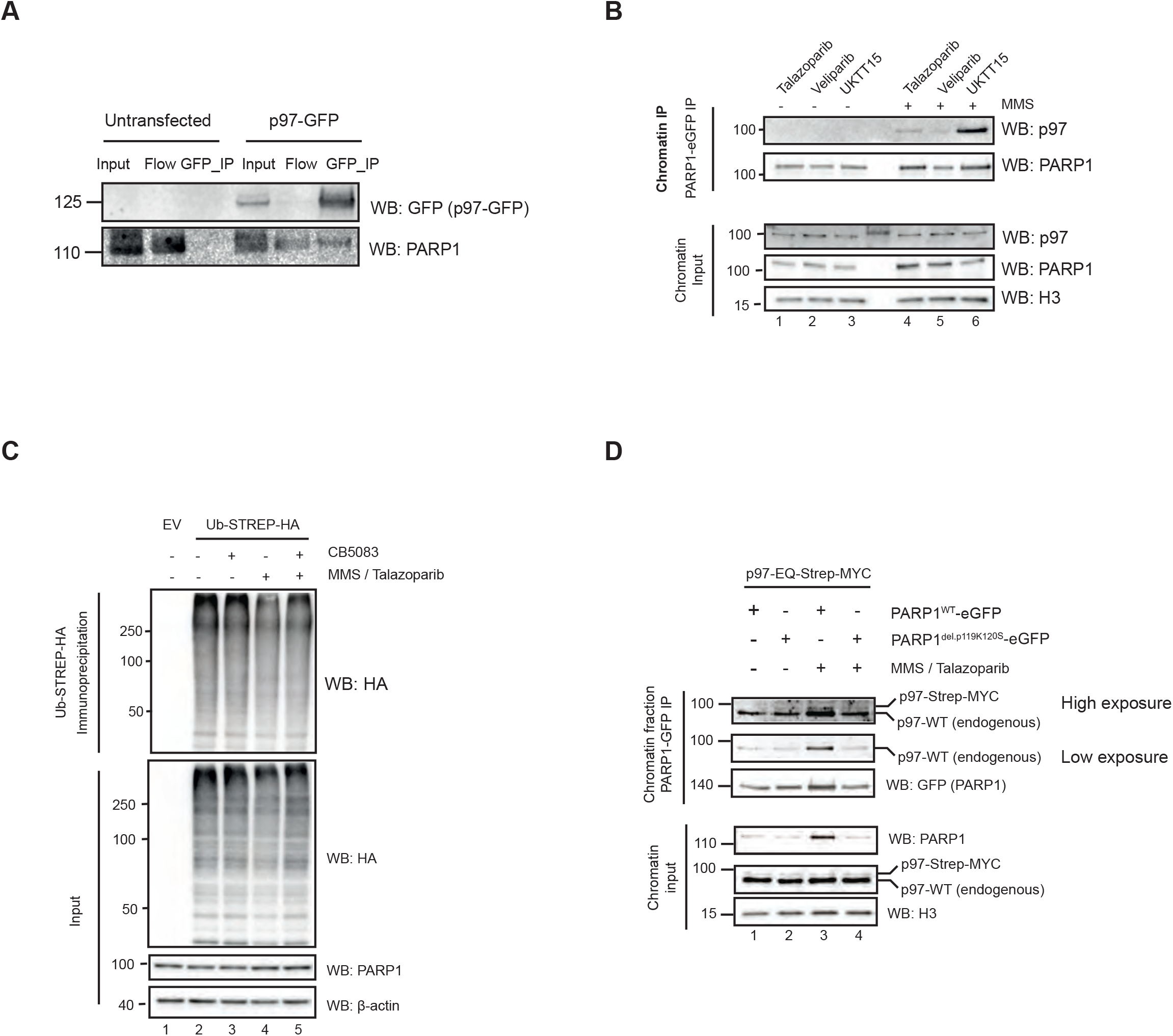
**A.** Western blot analysis of Co-IP confirms PARP1-p97 interaction. CAL51 cells were transiently transfected with p97-WT-GFP-expressing construct. Subsequently, GFP was immunoprecipitated in native conditions and the presence of PARP1 investigated by Western blotting. **B.** PARP1 interacts with p97 in a trapping-dependant manner. Cells were treated with 0.01 % MMS in the presence of 100 nM talazoparib, 10 µM veliparib or 10 µM UKTT15. PARP1 associated proteins were immunoprecipitated and the presence of p97 was investigated by immunoblotting. Only the trapping inhibitors, talazoparib and UKTT15, but not the non-trapping inhibitor veliparib led to an increased in the interaction between PARP1 and p97. **C.** Western blots for denaturing IP experiment shown in Figure 4F. **D.** CAL51 PARP1^WT^-eGFP or PARP1^del.p.119K120S^-eGFP expressing cells were transfected with p97-EQ-Strep-MYC-expressing construct for 18 h (to prevent aggregation artefacts caused by the expression of the dominant negative p97). Subsequently they were treated with trapping conditions and fractionated into chromatin and soluble fraction. PARP1-eGFP was immunoprecipitated from the chromatin fraction by GFP-Trap beads and the association with p97 was investigated by Western blotting.

**Supplementary Figure 5.**
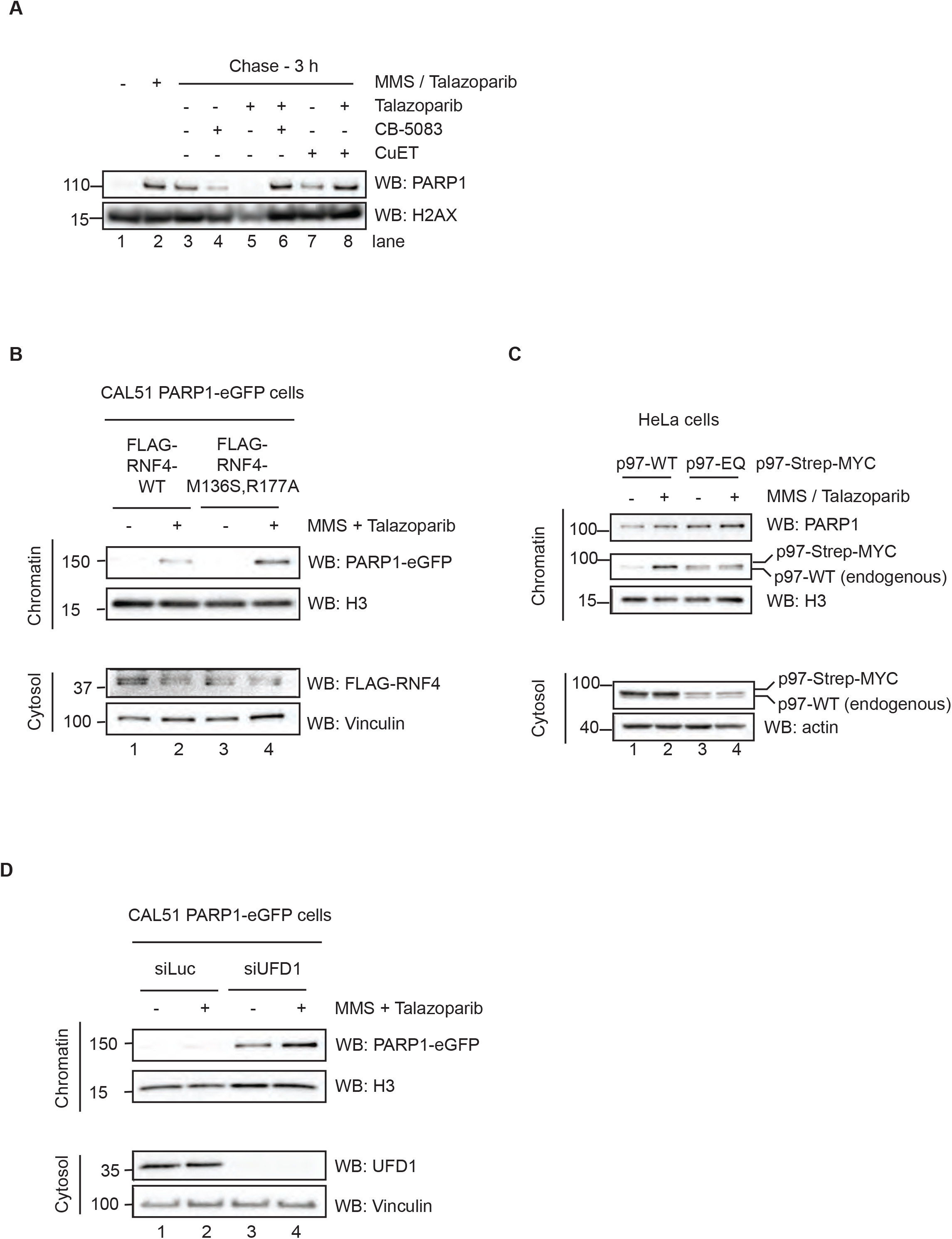
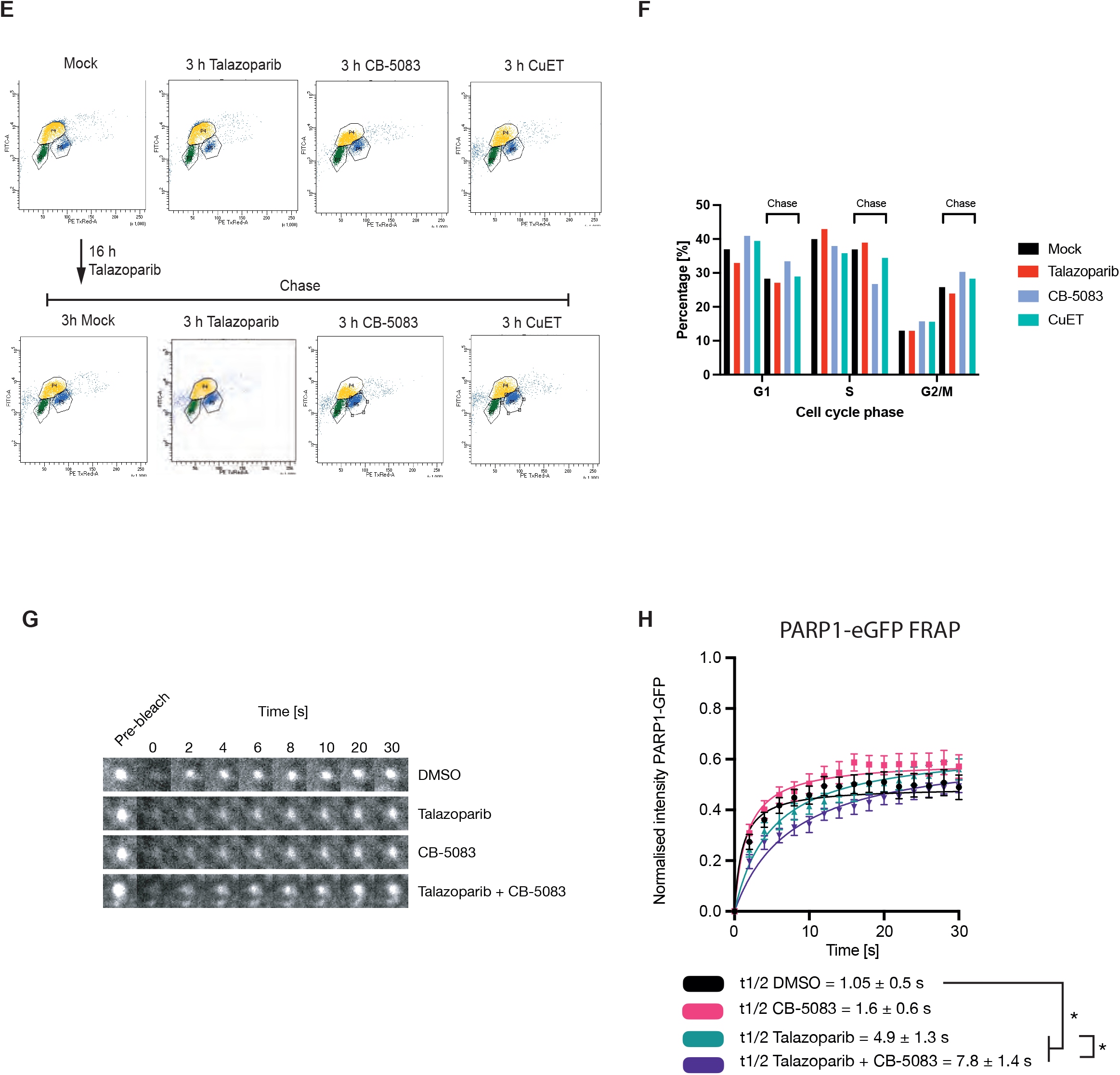
**A.** As described in Figure 5A, trapping was induced in cells and subsequently chased with media that was supplemented with PARPi (100 nM talazoparib) or p97 complex inhibitor (1 µM CuET or 10 µM CB-5083), either as single agent or in combination. At the end of the chase, the cellular nuclei were fractionated and the amount of chromatin-bound PARP1 was investigated by Western blotting. Trapped PARP1 was highest under the trapping conditions (lane 2), and decreased during the chase period in the single drug treatment arms (lanes 3, 4, 5 and 7); in contrast, the trapped complex persisted in the combinatorial arms (lane 6 and 8). Western blotting H2AX served as a loading control in these experiments. **B.** CAL51 PARP1^WT^-eGFP cells were transfected with FLAG-RNF4-WT or FLAG-RNF-M136S,R177A (E2 binding mutant, dominant negative) constructs. 24 h after expression the cells were treated with trapping conditions and subsequently fractionated into soluble and chromatin-bound fractions. The expression of the FLAG-RNF-M136S,R177A construct led to higher amount of trapped PARP1 in the chromatin fraction. **C.** HeLa cells were transfected with either p97-WT-Strep-MYC or p97-E578Q-Step-MYC (dominant negative) mutant expressing plasmid for 16 hours (to avoid the formation of overexpression artefacts) and subsequently treated with trapping conditions. The cells were fractionated into chromatin and cytosol and the accumulation of PARP1 in the chromatin fraction was investigated via Western blotting. Notably, although p97-E578Q mutant was expressed at a lower level compared to the p97-WT construct, it led to higher accumulation of PARP1 in the chromatin fraction. **D.** UFD1 is necessary for the efficient extraction of trapped PARP1 from the chromatin. CAL51 PARP1-eGFP expressing cells were transfected with a control siRNA (siLuc) or UFD1-targeting siRNA (siUFD1). 48 hours post transfection the cells were treated with trapping conditions and subsequently fractionated into cytosolic and chromatin-bound fraction. UFD1 depletion led to higher amount of chromatin bound PARP1, which was further increased by trapping in MMS/Talazoparib conditions. **E.** A cell cycle profiling for the experiment shown in Figure 5I. CAL51 cells were treated with the corresponding drug conditions. One hour prior to fixation, 10 µM EdU was added to the media. The cells were collected, EdU was stained by a click reaction with Alexa488-azide and DNA was stained by propidium iodide. **F**. A quantification of the G1, S and G2 populations from (D). The 3 h drug treatment arms do not alter the distribution of the cell cycle phases. In contrast, the 16 h Talazoparib treatment leads to an increase in the G2 population. This accumulation is not changed by the p97 inhibition treatment arms for the chase period of 3 h. **G.** CAL51 PARP1^WT^-eGFP cells subjected to microirradiation, where the accumulation of PARP1 at UV-laser induced DNA damage sites was monitored over time. At the maximum time of recruitment (typically 1 min after microirradiation) the focus was bleached with a 488 nm laser and the recovery of the GFP signal was monitored over time, essentially as described in *Shao Z. et al NAR 2020*. Shown are montages of the microirradiation site over time for the corresponding drug treatment arms. **H.** A quantification of the FRAP described in (F). The analysis was conducted as described in *Shao Z. et al NAR 2020*. The fluorescent signal was normalised as the maximum of recruitment immediately prior the photobleach to 1 and the signal immediately after the photobleach to 0. The recovery data was fitted with one site-specific binding model of non-linear regression and the extra sum of squares *F* test was used to calculate the *t_1/2_*. The significance was determined with a two-sided *t*-test from two independent experiments, where 10 to 12 cells were quantified for each condition. * - *p*-value < 0.05

**Supplementary Figure 6.**
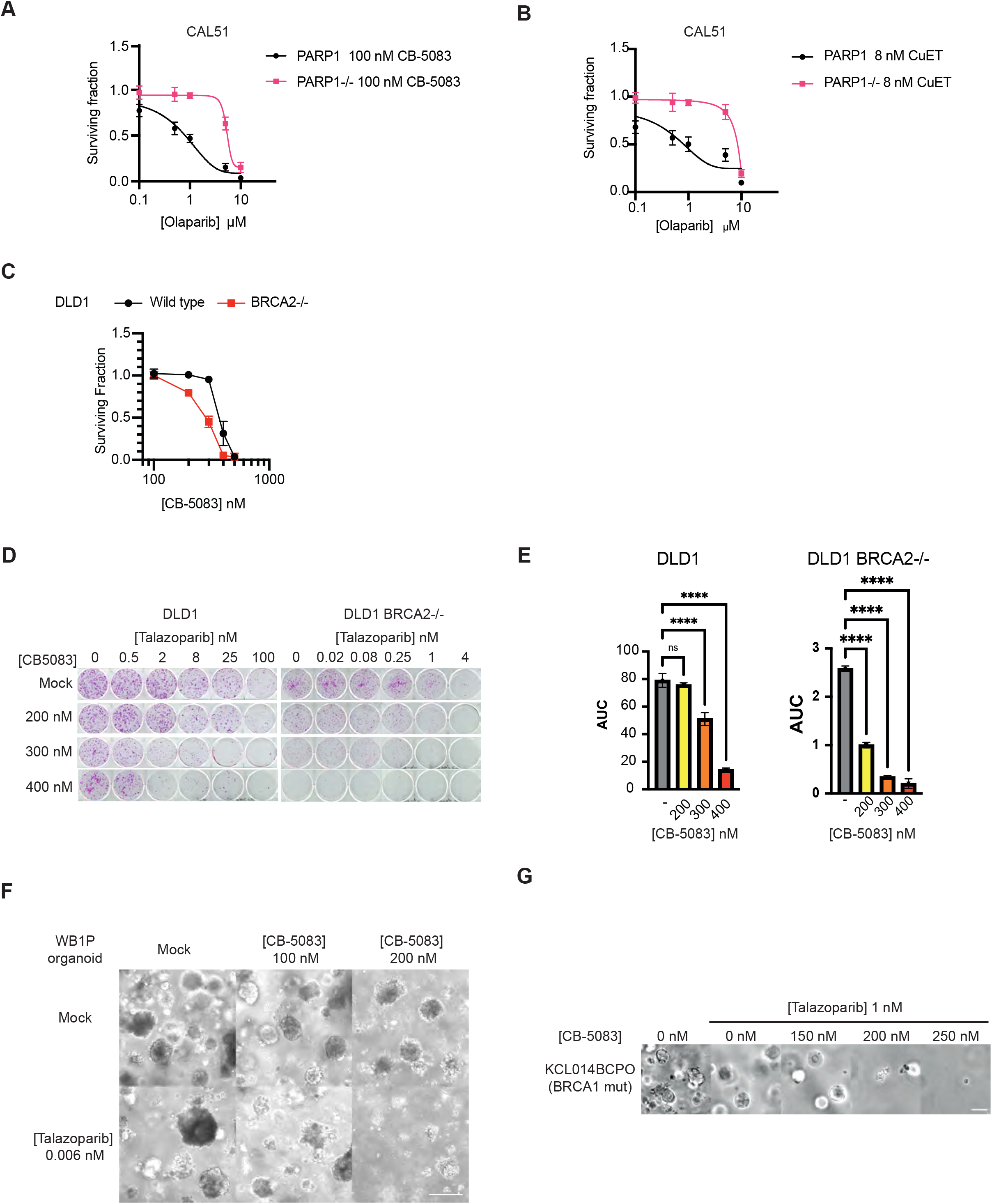
**A.** and **B.** Drug response curves for the colony formation assays presented in Figure 6A. CAL51 WT or *PARP1^-/-^* cells were treated with increasing concentrations of the PARP inhibitor Olaparib in the presence of either 100 nM CB-5083 (A) or 8 nM CuET (B). Surviving fractions were calculated based on the number of colonies after 14 days of exposure to the drugs. **C.** A quantification of the CB-5083 single agent effect on the surviving fraction of DLD1 and DLD1 BRCA2^-/-^ cells, respectively. **D.** Colony formation assays showing the synergistic effect between talazoparib and CB-5083 in DLD1 and DLD1 BRCA2^-/-^ cellular models. Increasing concentration of CB-5083 caused increasing sensitization to lower talazoparib concentrations in both models. Of note, the DLD1 BRCA2^-/-^ was treated with lower talazoparib concentrations as they are much more sensitive to PARPi. **E.** Area under the curve (AUC) analysis of the surviving fractions of DLD1 and DLD1 BRCA2^-/-^ cells in the presence of increasing concentrations CB-5083 combination as presented in Figure 6F. **F.** Brightfield images, showing the effect of talazoparib-CB-5083 combination on the GEMM WB1P organoid as described in Figure 6G. Scale bar represents 200 µm. **G.** Brightfield images showing the effect of talazoparib-CB-5083 combination on the KCL014BCPO organoid as described in Figure 6H. Scale bar represents 200 µm.

**Supplementary Table 1.** A list of proteins identified in the PARP1^WT^-eGFP RIME experiments.

**Supplementary Table 2.** A list of proteins identified in the *PARP1^del.p.119K120S^-eGFP* RIME experiments.

**Supplementary Table 3.** A list of proteins identified in the PARP1-Apex2-eGFP proximity labelling experiments.

**Supplementary Table 4.** Gene ontology terms for the proteins identified in the PARP1-Apex2-eGFP proximity labelling experiments.

